# Pre-saccadic suppression and compensatory cortical distortions support predictive visual remapping

**DOI:** 10.64898/2026.10.20.760565

**Authors:** Xize Xu, Sachira Denagamage, Anirvan S. Nandy, Monika P. Jadi

## Abstract

Perceptual continuity across saccades relies on the predictive remapping of receptive fields (RFs) of visual neurons, driven by corollary discharge (CD). However, how the brain achieves accurate and coherent remapping within its non-uniform visual representation remains unclear. Here, we present a generalized framework of recurrent 2D network in which neurons conjunctively tuned to retinotopic location and planned saccade direction integrate CD signals to predictively remap. We show that pre-saccadic suppression (PS) stabilizes this process by counteracting direction selectivity distortions, while recurrent connectivity preserves cell-cell RF relationship against input noise by constraining the cortical map to a low-dimensional manifold. Analyzing macaque V2 recordings, we validate saccade direction selectivity, the role of PS, and conserved RF relationships. Lastly, we show that while non-uniform representation introduces eccentricity-dependent remapping errors in the model, countervailing distortions systematically reduce them. Our study reveals how network architecture can reconcile the demands of visual acuity and perceptual continuity.

## INTRODUCTION

As animals interact with the environment, anticipating the sensory consequences of self-generated actions is essential for maintaining perceptual stability, and hence effective behavior. A key feature of the neural mechanisms supporting this anticipation is the preemptive modulation of activity in relevant brain areas prior to the action itself (1-5). The visual system provides a compelling example: ballistic eye movements known as saccades are a ubiquitous phenomenon that allows our visual system to examine objects of behavioral relevance in the visual scene. Each saccade induces a rapid displacement of the retinal image, thereby altering the visual information delivered to cortical neurons. Nevertheless, our perception of the visual world remains stable across saccades. One mechanism that enables this stability is predictive receptive field (RF) remapping (4, 6, 7), a phenomenon where neurons transiently shift the location of their spatial sensitivity before the onset of a saccade to compensate for the impending disruptions of visual inputs. Predictive remapping has been robustly observed in multiple visual and oculomotor areas, including V1, V2, V3, V3A, V4, lateral intraparietal cortex (LIP), frontal eye field (FEF), and superior colliculus (SC) (7-19). The advance information about the trajectory of the upcoming eye movement that is crucial for predictive remapping, has been shown to be conveyed by a corollary discharge (CD) signal originating from brain regions responsible for generating eye movements (4, 6, 20).

Maintaining perceptual stability requires high-quality remapping across two distinct levels. At the single-neuron level, the RF should remap precisely along the saccade vector to provide an accurate prediction of the post-saccadic visual frame (Fig. 1A2). At the network level, population remapping should remain coherent: conserved cell-cell RF relationships (Fig. 1A4), and a stable cortical population RF map constrained to a low-dimensional retinotopic space (‘manifold’). The quality of predictive remapping in the cortex, at either single neuron or population level, is not well understood. Additionally, factors that determine the quality of remapping remain unexplored.

**Figure 1.**
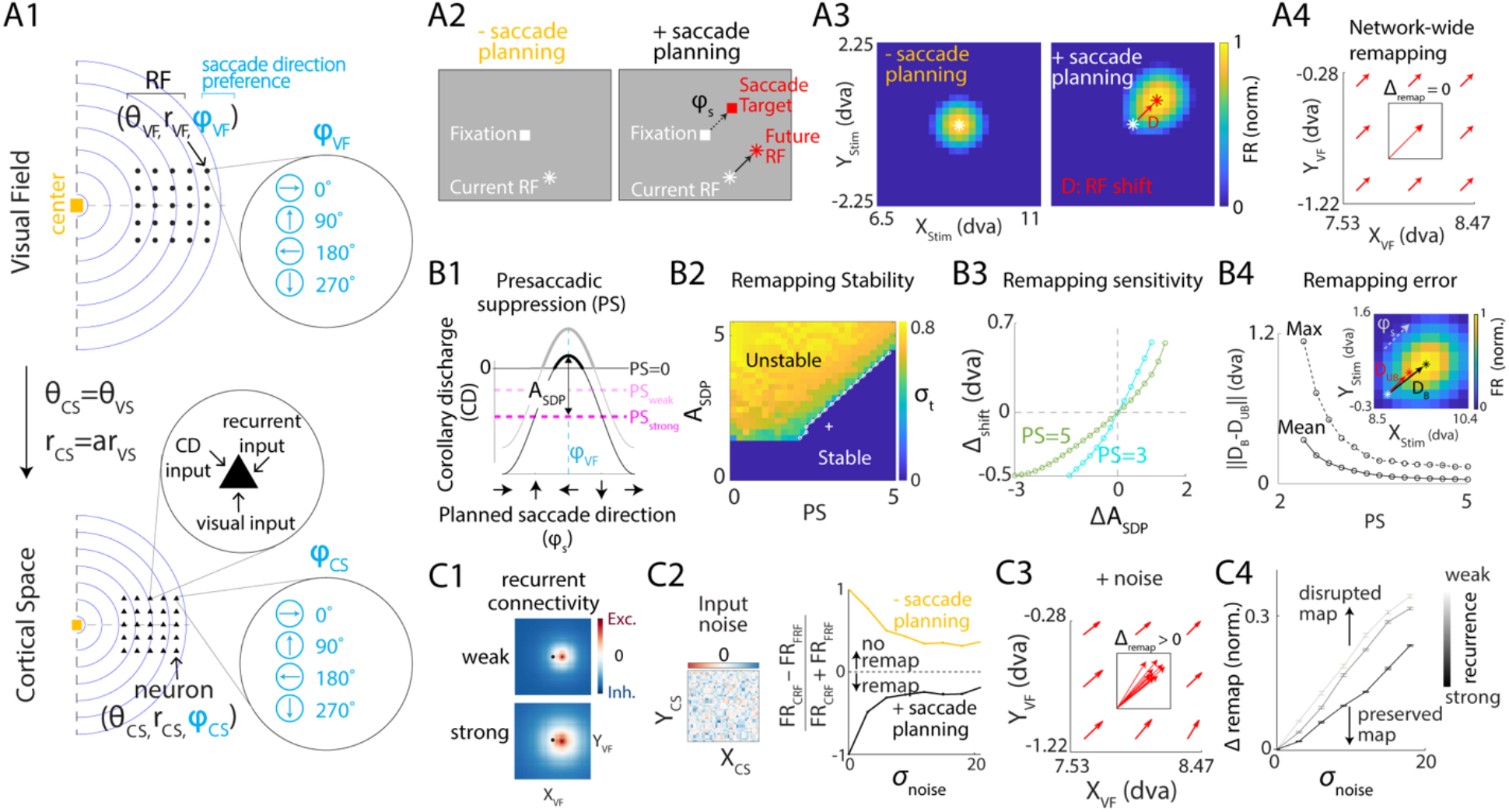
Network model of receptive field (RF) remapping under uniform cortical transformation. (**A**) Model description. **A1:** Schematic of the network modeling a patch of the cortical sheet. Each neuron in the network (filled black triangle) is assigned a spatial RF (filled black circle) and a saccade direction preference (SDP) in visual space. Iso-radius contours reflect uniform mapping between the cortical sheet and visual space. Dashed lines represent the meridians and their cortical representation. Each neuron receives recurrent input, external visual input and corollary discharge (CD) input. **A2:** Schematic of the experimental phenomenon of RF remapping. Left: RF of an example neuron during fixation without saccade planning, defined as current RF (CRF). Right: Under saccade planning, the RF predictively shifts by the planned saccade vector (φ_S_ ), defined as future RF (FRF). **A3**: Steady-state response (referred to as ‘ratemap’) of model neurons with identical spatial RFs to visual stimuli presented at different spatial locations during saccade planning. White, red asterisks: RF locations determined by the centroids of the ratemaps without (left) and with (right) saccade planning, respectively. dva: degrees of visual angle. **A4:** RF shifts of neurons under saccade planning sampled across cortical sites. Inset: RF shifts vectors aligned by their origins. (**B**) Role of pre-saccadic suppression (PS) in RF remapping. **B1:** Corollary discharge input as a function of planned saccade direction. Schematic is shown for neurons with example SDP (blue dashed line). Various levels of PS are implemented as a negative shift in the baseline (magenta lines). Thicker portions of curves highlight excitatory CD inputs for various levels of PS. A_SDP_: Amplitude of SDP. **B2:** Effect of PS on remapping stability. The heatmap illustrates standard deviation (σ_t_) of the temporal fluctuations in population activity centroid during saccade planning as a function of pre-saccadic suppression and amplitude of SDP. White line: boundary separating regimes of stable remapping (σ_t_ = 0) and unstable population activity (σ_t_ > 0). White plus sign: parameter values used in the example shown in **A3. B3:** Effect of PS on the sensitivity of remapping to SDP. We assessed the robustness by quantifying the shift in stationary network activity bump as a neuron-averaged measure of RF remapping of individual neurons (see Fig. S1C). Curves show variation in the planning-induced shift (Δ_shift_ ) of stable activity centroid (Fig. S1C), as a function of deviation in A_SDP_ (Δ A_SDP_). PS and A_SDP_ values at baseline (0,0) yielded the same shift (A_SDP_ = 1.4 for PS = 3, A_SDP_ = 2.6 for PS = 5). **B4:** Effect of PS on the remapping error caused by non-uniform SDP. Maximum and average remapping error due to SDP amplitude bias (Fig. S1D1) as a function of PS level, across all tested planned saccade directions (Fig. S1D2). Inset: Remapped RF of neurons with identical CRFs (white asterisk) during saccade planning, but in networks with (black asterisk) and without (red asterisk) biased SDP (Fig. S1D1). The ratemap in background shows activity during saccade planning in a network with biased SDPs. (**C**) Effect of recurrent connectivity on cell-cell RF remapping coordination in the presence of noise. **C1:** Illustration of weak and strong recurrent connectivity as seen in outgoing weights from neurons with an exemplar spatial RF (black dot) and SDP (same as in B1) to postsynaptic neurons located across visual field. **C2:** Left: Illustration of independent input noise across cortical locations. Right: Remapping sensitivity index (RSI) as a function of noise level (σ_noise_ ) without (yellow) and with (black) saccade planning. 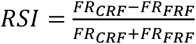, where FR represents the neuronal response at current RF and FR_,C,_ represents the response at the expected forward remapped RF. A negative RSI indicated remapping. **C3:** Same as A4 but in the presence of input noise. **C4:** Normalized pairwise difference in RF shifts (normalized 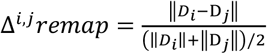) as a function of the standard deviation of noise (σ_noise_) across various levels of recurrent connectivity strength (Methods). Error bars indicate the 95% confidence interval for the mean.

One factor that can influence remapping quality is a co-occurring phenomenon called pre-saccadic suppression (PS). It is a temporal reduction in neuronal firing rates during saccade planning (9, 21-25). Empirical studies suggest a correlation between extent of PS and remapping. For example, V1 neurons show limited suppression or even facilitation of firing rates (4, 26), compared to those in V2 or V4 (9, 21), during visually guided saccade tasks . This parallels V1 neurons showing weaker remapping compared to higher-order visual areas, including V2 and V4 (8, 9, 19). These observations suggest the possibility that PS may play a functional role in remapping.

A factor that can influence the remapping quality at the population level is the strength of recurrent connectivity (27-29), especially in the presence of uncorrelated noisy inputs during remapping. While recurrent connectivity is expected to be crucial for remapping (30, 31), how it influences the relaxation to this low-dimensional manifold, is not known.

Most importantly, a factor that is fundamental to visual cortical organization but remains completely unexplored is the impact of non-uniform cortical transformations on remapping quality. In a foveated visual system, more neural resources are dedicated to processing visual information located at or near the fovea than at peripheral locations, resulting in eccentricity-dependent cortical magnification (CM). Both CM and RF size change with eccentricity (32-36), thus resulting in a non-uniform representation of visual space. This eccentricity dependence increases along the ventral visual pathway (34, 37-39). Additionally, the representation of polar angles is contracted over a substantial region surrounding the fovea in the transformation from the visual field to cortical space (37, 38, 40, 41). It is not known how these non-uniform geometric characteristics interact and impact the accuracy of predictive remapping.

Both RF remapping errors and non-uniform transformations occur in the 2-D visual and cortical spaces, respectively. Thus, addressing these questions requires a unified approach that integrates 2-D computational modeling with validation in 2-D empirical data to provide deeper insights into the quality of remapping. Further, visual areas differing both in the extents of PS and non-uniform representations demand a generalized framework that can explore this variability.

In this study, we present a generalized architectural framework of 2-D cortical space to model forward remapping, wherein RFs shift along the planned saccade vector. While CD inputs predictively remap RFs via a network mechanism involving neurons conjunctively selective to planned saccade direction and visual field location, the framework allows us to systematically explore the role of the factors that can shape the quality of remapping.

We reveal a regulatory role for PS to counteract the adverse effects of saccade direction selectivity during saccade planning, a prediction validated by reanalyzing a high-density dataset in macaque V2. Our model reveals competing effects of input noise and recurrent connectivity on cell-cell RF relationships. Cortical transformations such as eccentricity-dependent CM and polar angle contraction systematically distort remapping in our model, findings that are confirmed by the empirical data. Finally, our model proposes that the differential distortion of the cortical map by each type of transformation can cause a mutually mitigating effect, thus reducing predictive remapping error.

## RESULTS

### Network model of receptive field remapping under uniform cortical transformation

To investigate the computational mechanisms underlying predictive remapping, we modeled the visual cortex using a dynamical rate network (42-44) arranged on a 2-D cortical sheet representing a small part of the visual field. Given minimal receptive field (RF) transformations at a small spatial scale, we assumed the neurons’ cortical location to be uniformly scaled from the location of their RFs (Fig. 1A1). As a result, the overall shape that bounded a set of RF locations was preserved when mapped onto the cortical sheet. The model neurons were conjunctively selective to visual field location and planned saccade direction. Thus, each possible RF location is represented by a group of neurons with distinct saccadic direction preferences (SDPs; Fig. 1A1). Unless otherwise indicated, we assumed uniform strength of saccade direction selectivity across neurons with different SDPs. The outgoing connection weight from each neuron consisted of a fixed level of inhibition and an excitatory component that decays exponentially with the distance from target neurons, with the profile center shifted opposite to its SDP (Fig. S1A; Methods). Inhibitory connections between neurons with distant RFs stabilized network activity by preventing runaway excitation. Visual stimulus was modeled as an external input whose amplitude decreased as a function of the distance between the representation of the visual stimulus and a neuron’s cortical location (Fig. S1B; Methods). To simulate saccade planning, an additional input modeled the corollary discharge (CD) associated with a planned saccade. Its amplitude depended on the level of pre-saccadic suppression (PS) and was modulated by the alignment between the planned saccade direction and the SDP of each neuron (Fig. 1B1; Methods). PS was incorporated into the CD, rather than any visual or proprioceptive input, based on empirical evidence that it occurs even in the absence of visual stimulation, or with paralyzed eye muscles (see (4) for a review). The network operated in a dynamical regime with no self-sustained activity, remaining silent in the absence of external inputs. In the absence of saccade planning, but with visual stimulation, the network stabilized to a stationary population activity bump representing the visual stimulus (Fig. S1C, left). Under saccade planning, neurons with the same RFs but distinct SDPs were differentially activated by the CD inputs, creating an imbalance that propagated the population activity opposite to the planned saccade direction. When combined with a visual stimulus, the CD input could lead to a stationary population activity bump shifted away from the representation of the visual stimulus location, in a direction opposite to that of the planned saccade (Fig. S1C, right), thus changing the perceived location of the visual stimulus (45). Effectively, the model captured the empirical phenomenon of predictive RF remapping (Fig. 1A2) as the shift in stationary population activity bump translated to the remapping of current RFs (CRF) of individual neurons along the saccade direction (Fig. 1A3; see Methods). At the population level, as the connectivity profiles and receptive inputs were invariant across cortical sites, the RF shifts were uniform across the network, implying a coherent spatial representation despite substantial changes in single-cell RFs (Fig. 1A4).

### Pre-saccadic suppression enhances stability, robustness and accuracy of predictive remapping

Inherent temporal variability in the biological recurrent circuit properties can degrade remapping performance by introducing instability or inaccuracy. For a stable percept, it is thus critical for the network to remain robust to such variability. Saccade planning has been shown to be accompanied by a temporary reduction in neuronal firing rates across various visual cortical areas (9, 21-23), a phenomenon known as ‘pre-saccadic suppression’ (PS). Given its prevalence across visual areas that perform predictive remapping, and its temporal overlap with the remapping process (see Discussion), we reasoned that PS might act as a physiological counterweight to temporal circuit variability.

By incorporating PS as modulation of the CD input in our model (Fig. 1B1), we first probed its impact on the network’s ability to perform stable remapping. Specifically, we varied the amplitude of saccade direction selectivity (*A*_*SDP*_) and measured the standard deviation of the population activity centroid (*σ*_*t*_) during saccade planning, as a function of PS. Increasing *A*_*SDP*_ required a stronger PS to obtain stable remapping (*σ*_*t*_ = 0) in the model network. Thus, the range of *A*_*SDP*_ allowing stable remapping became broader with stronger PS (Fig. 1B2). These results suggest that PS regulates the destabilizing effect of *A*_*SDP*_, keeping the network in a stable remapping regime.

We next examined the impact of PS on the robustness of remapping under varying values of *A*_*SDP*_ . Given the functional coupling between RF remapping of individual neurons and the shift in stationary network activity bump in our model, we assessed the robustness by quantifying the latter as a neuron-averaged measure. For each level of PS, we selected a reference value for *A*_*SDP*_ so as to maintain a constant shift in the population activity centroid under steady state during saccade planning (Fig. S1C, right). We then determined how this shift varied with the deviation in *A*_*SDP*_ from the reference value within the regime of stable remapping (Fig. 1B2). The shift of the population activity centroid was less sensitive to changes in *A*_*SDP*_ for greater PS (Fig. 1B3), implying a role for PS in providing robustness to temporal variability in *A*_*SDP*_.

Beyond stability and robustness, we investigated the impact of PS on remapping accuracy. While multiple factors can generate remapping errors, we focused on the bias in the distribution of SDPs within visual areas as suggested by recent experimental results (46, 47) and tested how PS modulates the resulting remapping errors. We simulated this bias by increasing the *A*_*SDP*_ for neurons with a specific SDP (Fig. S1D1). Compared to the unbiased case in which the RF shift consistently followed the direction of the planned saccade and was thus considered perfect remapping, non-uniform SDP resulted in biased remapping (Fig. 1B4, inset, Fig. S1D2). We characterized the error induced by this bias as the magnitude of the difference between the locations of remapped RFs with and without bias. Stronger PS attenuated the remapping errors induced by the bias (Fig. 1B4), suggesting that PS can also help maintain remapping accuracy despite biases in SDPs.

Collectively, these results suggest that while saccade direction selectivity is essential for predictive remapping, it is also a source of instability, variability, and error in remapping. Crucially, PS serves as a counteractive mechanism, mitigating these adverse effects and maintaining stability, robustness, and accuracy during remapping.

### Recurrent connectivity improves the robustness of cell-cell RF coordination to input noise during remapping

Although recurrent connectivity is a prominent feature of visual cortex (27-29), its functional role in maintaining population-level spatial coherence under neuronal noise — particularly across saccades involving displacement of the visual frame, rather than during steady-state fixation — remains to be explored. At the population level, predictive remapping can occur along a spectrum: from an uncoordinated regime, where single-cell RFs shift independently, to a coordinated regime, where spatial relationships between cell RFs are conserved despite single-cell RF shifts. The latter regime implies a spatially coherent representation across saccades.

To address this question, we first examined the impact of external input noise on the population dynamics of remapping. Uncorrelated Gaussian input noise was injected into each neuron (Fig. 1C2, left; Methods), with its level characterized by the standard deviation *σ*_*noise*_ . We varied the noise level ( *σ*_*noise*_ ) and characterized the extent of coordination across neurons during remapping via the mean pairwise difference in RF shifts ( ||*Δremap*|| ; Methods). Neurons were perfectly coordinated during remapping without noise (Fig. 1A4), whereas they exhibited variance in RF shifts under noise (Fig. 1C3), with its amplitude increasing as a function of the noise level (Fig. 1C4). Crucially, during this disruption of RF relationships, individual neurons still satisfied classic experimental criteria for single-unit remapping (9), evidenced by stronger responses to the putative ideal remapped RF along the saccade direction ( *φ*_*S*_ ) (Fig. 1C2, right; Methods). Having established that input noise degrades spatial coordination, we systematically varied connection strength (Fig. 1C1; Methods) and found that the noise-induced disruption in population RF relationships could be rescued by stronger recurrent connectivity (Fig. 1C4). These results suggest that recurrent connectivity buffers the network against input noise during remapping, allowing it to preserve cell-cell RF relationships and thereby maintain a relatively coherent representation.

### Visual cortical activity during saccade planning

To test key model assumptions and predictions, we reanalyzed a dataset from our group that recorded the spiking activity of simultaneously recorded and well-isolated single neurons within cortical columns of visual area V2 of two awake macaque monkeys performing a cued-saccade task (9) (Fig. 2A-B). The subjects were required to acquire and maintain fixation (Fig. 2B, yellow square) for a variable delay period (500-900 ms, drawn from an exponential distribution), execute a saccade in response to the appearance of a target point (Fig. 2B, blue squares) in the periphery, and then continue holding fixation at the target point to receive a reward (Fig. 2C). The target location was drawn from one of two possible locations equidistant from the fixation point, cueing saccades with directions 45° apart from the axis connecting the receptive field of the recorded column and the fixation point (Fig. 2B; Methods). Both the saccade target location and the delay period duration were pseudo-randomized to prevent preemptive planning of a saccade prior to the go cue. While the subjects executed these eye movements, oriented Gabor stimuli were continuously presented at 60 Hz on a 13 x 13 grid spanning the visual region of interest (Fig. 2B; Methods). On each frame of stimulus presentation, a single stimulus drawn from one of six random orientations was presented at a single grid location. The period from the onset of stimuli presentation, which was 200 ms after fixation acquisition, to the appearance of the go cue is referred to as the ‘pre-planning period’ (Fig. 2C). The location and orientation of the stimulus grid were determined by the receptive field of the recorded column on a session-by-session basis (Methods) to ensure that the relationship between fixation point, saccade target locations, and receptive field locations was conserved across sessions. In total, 923 single units were recorded, 377 of which exhibited substantial spiking responses (> 3Hz), had significant receptive fields (Methods), and were included in our subsequent analysis. Previous work has shown that the recorded neurons underwent remapping during the task across all cortical layers (9) (Table S1). While this dataset was previously used for remapping identification based on the neuronal responses at current and future RFs alone, here we leverage the full spatial profile of these recordings to precisely characterize RF locations and their temporal evolution across saccades, allowing us to directly test model predictions.

**Figure 2.**
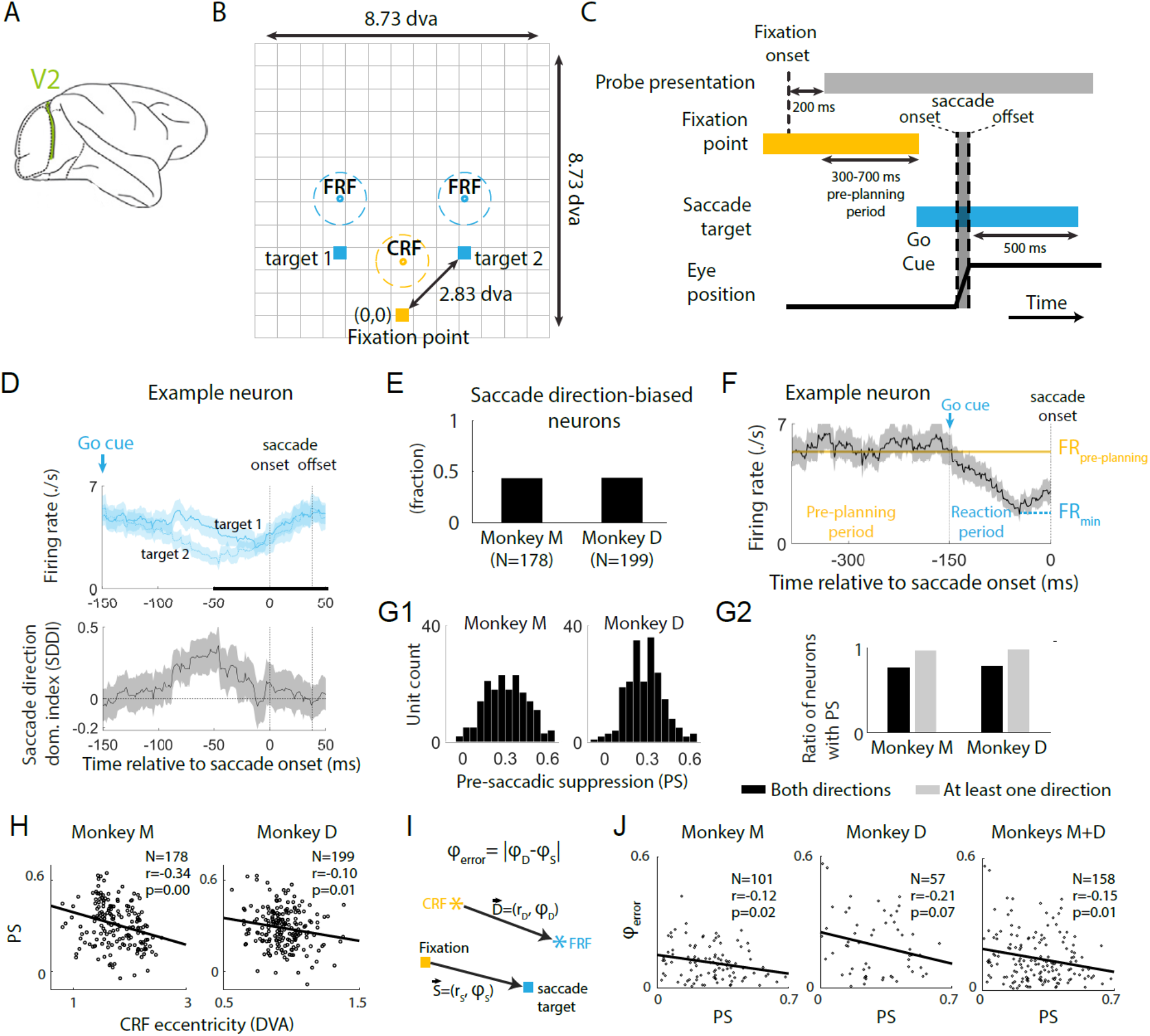
Characterization of planned saccade direction bias and pre-saccadic suppression in V2. (**A**) Illustration of recording locations. Populations of cortical area V2 units were simultaneously recorded with high-density laminar probes (2 shanks with 32 channels/shank) from two monkeys performing a cued saccade task. (**B**) Fixation point, saccade targets, and stimulus grid layout during the cued saccade task. (**C**) Schematic of the cued saccade task. Probe stimulus was presented at 60 Hz, with spatial locations randomized across the grid. (**D**) Example V2 neuron responses to the two saccade directions. Top: peri-event time histograms. Bottom: saccade direction dominance index (SDDI; Methods). Positive SDDIs indicate stronger responses to target 1, whereas negative SDDIs indicate stronger responses to target 2. Shaded areas indicate the 95% confidence interval for the mean. Bold portion of time axis indicates significant remapping period. (**E**) Ratio of recorded units exhibiting differential responses for saccade targets from each monkey. The results were calculated from activities in response to stimulus flashes presented during the significant remapping period. (**F**) Example V2 neuron that exhibits suppressed firing rate during the reaction period following target appearance compared to the pre-planning period. Horizontal line: baseline firing rate determined from the pre-planning period. (**G**) Pre-saccadic suppression (PS) across population. **G1**: The distribution of PS strength (Methods), with positive values indicating suppression. **G2**: Ratio of recorded units exhibiting significant PS. (**H**) PS strength averaged across saccade targets as a function of CRF eccentricity for each monkey (Methods). Black line: linear fit to the data. (**I**) Illustration of remapping direction error (φ_error_). (**J**) φ_error_ as a function of PS strength for each monkey and combined across monkeys (Methods). Black lines: linear fits.

### Saccade direction bias in V2

Our model predicted that saccade direction selectivity of individual neurons was essential for remapping by shifting the population activity opposite to the planned saccade direction (Fig. S1C). To test this model prediction, we compared the spiking responses of individual neurons during saccades cued towards targets in different directions (Fig. S2A; Methods). For a representative neuron, a difference in responses across saccade directions emerged after the appearance of saccade targets and, notably, before saccade onset (Fig. 2D, top). To quantify this difference in the spiking responses for each neuron, we introduced a saccade direction dominance index (SDDI). SDDI was significantly signed for a substantial time during saccade planning (Fig. 2D, bottom), indicating a bias in response towards one of the saccade directions. We found that about 43% of V2 neurons (Fig. 2E; 77 of 178 for monkey M, 87 of 199 for monkey D) exhibited significantly signed SDDI during the remapping period (-50 to 50 ms relative to saccade onset), and therefore had biased responses across the two saccade directions, suggesting saccade direction selectivity in V2 during saccade planning.

### Pre-saccadic suppression (PS) reduces remapping errors in V2

Our model predicted that PS enhanced the accuracy of remapping. To test this prediction, we first evaluated the presence of PS. For a representative neuron, the firing rate decreased significantly following the appearance of the saccade target, with this suppression emerging notably before saccade onset (Fig. 2F). To quantify this modulation of spiking activity during saccade planning, we calculated the relative change in firing rate during the pre-saccadic period compared to the pre-planning period prior to target appearance (Fig. 2F; Methods). We found that about 78% of V2 neurons exhibited a significant reduction in firing rate during saccade planning (137 of 178 for monkey M, 157 of 199 for monkey D) under both saccade directions, about 98% (173 of 178 for monkey M, 196 of 199 for monkey D) for at least one saccade direction (Fig. 2G), demonstrating robust pre-saccadic suppression across the population. Notably, we observed variability in PS strength across neurons (Fig. 2G1), which might reflect noise within corollary discharge signals during saccade planning. Moreover, to assess whether the strength of PS is uniform across the cortical sheet, we examined the relationship between PS strength and the eccentricity of the RF estimated before the appearance of the go cue, referred to as the ‘current receptive field’ (CRF). Interestingly, we found that pre-saccadic suppression systematically decreased with CRF eccentricity (Fig. 2H), suggesting a spatial gradient where suppression is stronger in neurons representing foveal visual space compared to those representing the periphery.

Having established the presence of PS, we next investigated the relationship between PS strength and remapping accuracy. For each neuron, we characterized the stable remapped RFs and the corresponding RF shifts during saccade planning (Methods), interpreting these as the steady-state responses of the network model to the combined influence of visual stimulus and CD inputs. Consistent with the model prediction, remapping direction errors, computed as the angular difference between RF shift direction (*φ*_*D*_) and saccade direction (*φ*_*S*_) (Fig. 2I), decreased with PS strength in both monkeys (Fig. 2J). This facilitating role of PS thus provides further support for the proposed theoretical framework.

### Cell-cell RF relationships remain relatively stable despite transient degradation during predictive remapping

We next characterized cell-cell RF relationships to test the model’s prediction regarding their sensitivity to input noise, as reflected by the variation in PS strength across neurons (Fig. 2G1). The continuous presentation of visual stimuli at various locations on the 13×13 grid across saccades allowed us to estimate the RFs for each neuron on a moment-by-moment basis. We binned stimulus flashes presented at each location with a 41 ms window according to their presentation times relative to saccade onset. The average responses to these flashes formed a 13×13 map of spike counts across stimulus locations for each neuron at each timepoint relative to saccade onset. Focusing on the visual region of interest (Fig. S2B, C), we estimated the RF of each neuron as the centroid of locations where stimuli caused sufficiently high spiking responses. The temporal evolution of the RF for each neuron also allowed us to estimate its shift during remapping, determined as the vector from the CRF to the temporal RF (Fig. 3A). We characterized the cell-cell RF relationship by calculating the pairwise difference in RF shifts (*Δremap*; Fig. 3B). Small amplitudes of *Δremap* indicate conserved RF relationships, whereas large amplitudes indicate disrupted RF relationships.

**Figure 3.**
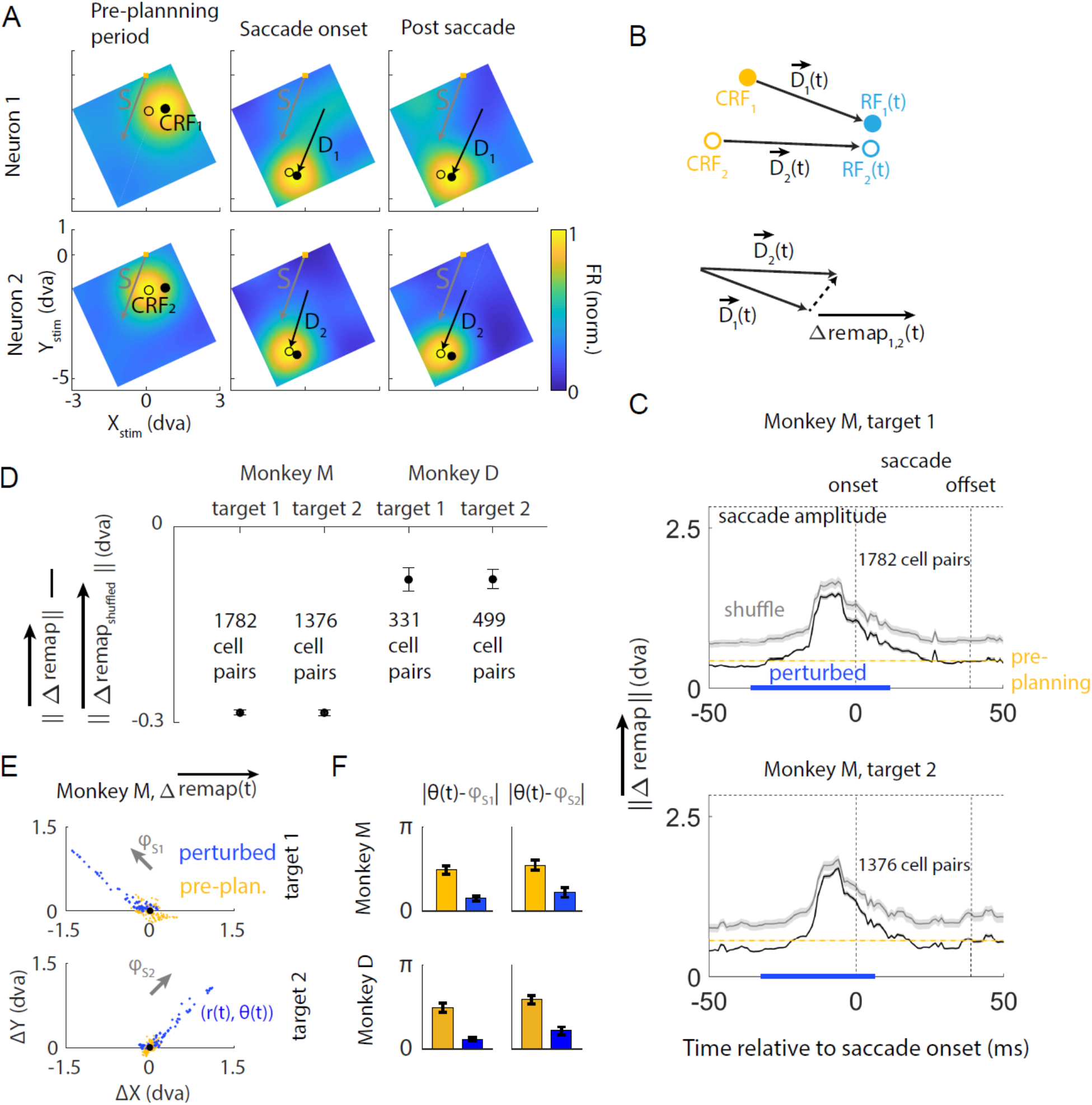
Characterization of the dynamics of cell-cell RF relationships in V2. (**A**) RF locations for an example pair of V2 neurons (from the same session and cortical layer) during the cued saccade task. Filled and open circles indicate the RF locations of neuron 1 and neuron 2, respectively. Yellow squares: fixation points. Grey arrows: saccade vectors. Black arrows: RF shifts from CRFs to temporal RFs. Heatmaps are individually normalized. (**B**) Illustration of pairwise difference in RF shifts (Δremap). (**C**) The temporal evolution of the mean 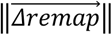 for neurons pooled across sessions from monkey M for each saccade target (black). 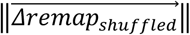 was estimated using shuffled CRFs (grey). Only RFs of neurons with the same laminar identities (Methods) were paired or shuffled. Gray dashed line: saccade magnitude / expected RF shifts for individual neurons. Yellow dashed line: mean 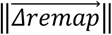 during the pre-planning period. Blue bar: period during which 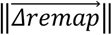 was persistently higher than the pre-planning level, termed the ‘perturbed period’. Shaded areas indicate the 95% confidence interval for the mean. (**D**) Difference between mean 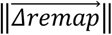 and 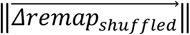 for each monkey. Error bars indicate the 95% confidence interval for the mean. (**E**) Deformation of Δremap during the perturbed period. Each dot represents the mean Δremap at one timepoint (advancing by 1 ms) during the perturbed period (blue) or the pre-planning period (yellow) across all cell pairs pooled across sessions for monkey M. The temporal average (black circle) during the pre-planning period was taken as zero. Grey arrows: saccade directions (φ_S_). (**F**) Absolute difference between deformation direction (θ(t)) and saccade direction (φ_S_).

Assuming rotation-invariance, we combined the neural data across sessions by aligning the session-specific stimulus grids. All RFs and their shifts were calculated in the reference frame of the stimulus grid, where the coordinates of the fixation points as well as saccade targets, and thus the saccade directions, were fixed. The mean 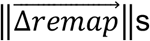 remained low until about 25ms before saccade onset, then increased and dropped back to the pre-planning level (Fig. 3C) before saccade completion, after which the feedforward retinal inputs were realigned to the shifted visual frame. Interestingly, this transient increase, which reflected a degradation of cell-cell RF relationships, temporally overlapped with the window in which PS emerged during saccade planning (see Discussion). Given the substantial variability in PS strength across neurons (Fig. 2G1), this observation suggests a degrading effect of input noise on the coordination of remapping, in alignment with our model prediction. Despite the transient rise, throughout the remapping process, the 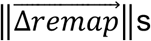 were consistently lower than the RF shifts of individual neurons, given by the amplitudes of cued saccades (Fig. 3C). This implies that the RF relationships between neurons were more stable than the RFs of individual neurons. Furthermore, the 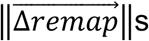 4s were lower than those calculated using shuffled CRFs (Fig. 3C), with a smaller difference for one subject—possibly due to the tighter distribution of the CRFs of recorded neurons (Fig. S2D, E). This result was robust across saccade targets for each monkey (Fig. 3D).

### Cell-cell RF relationships deform along the saccade vector

Since the CD input encodes the upcoming saccade, we further investigated whether it transiently modulated the cell-cell RF relationships in a systematic way. For each cued saccade direction, we computed *Δremap*s during the ‘perturbed period’, defined as the period during which 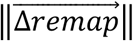 was persistently higher than the pre-planning level (Fig. 3C). The difference between the *Δremap*s during the perturbed and pre-planning periods characterized the perturbation of the original cell-cell RF relationships (defined by retinotopy) due to saccade planning (Fig. 3E). For both cued saccade directions, the transient deviations in *Δremap* during the perturbed period were aligned with the planned saccade directions (Fig. 3E, F), suggesting that these shifts were indeed induced by CD inputs.

### Network model of RF remapping under non-uniform cortical transformation

The representation of the visual field in visual areas is not uniform due to eccentricity-dependent cortical magnification (CM) (32-36) and the contracted representation of polar angles (37, 38, 40, 41). We incorporated these non-uniformities into our model to investigate their impact on the quality of predictive remapping across the visual field and cortical space. Since RF remapping is a robust phenomenon observed across multiple stages of the ventral visual stream, we sought to build a generalized architectural framework rather than one restricted to a single cortical area. In our model, a location in the visual field *(r*_*VF*_, *θ*_*VF*_) was mapped to a cortical location (*r*_*CS*_, *θ*_*CS*_) via a logarithmic transformation of its radius (*r*_*VF*_) and a contraction of its polar angle (*θ*_*VF*_) by a factor *k* (Fig. 4A). The logarithmic CM was achieved by implementing a linear relationship between RF size and eccentricity (48) (Methods). To examine how cortical magnification affects remapping quality, we explored a range of the slope (*s*) of this RF size-eccentricity relationship (37, 38) based on previous experimental measurements, which varied across different stages along the ventral stream. While all nonzero slope values produce logarithmic CM, indicating identical eccentricity dependence of CM (subject to scaling; Fig. S5C), a larger *s* value induces greater cortical distance-dependence of CM, marked by a faster decay rate of CM factor (CMF) across cortical locations along an iso-polar contour (Fig. S5D; Methods). With polar angle contraction (*k* < 1, quantifies the extent of contraction), a hemifield in visual space (center angle = π) is represented in cortical space in the shape of a minor sector (center angle = *k*π )(Fig. 4A). For modeling convenience, the SDPs of a neuron were restricted to the radial as well as its orthogonal and opposite directions, as defined in the visual field (Fig. 4A; Methods).

**Figure 4.**
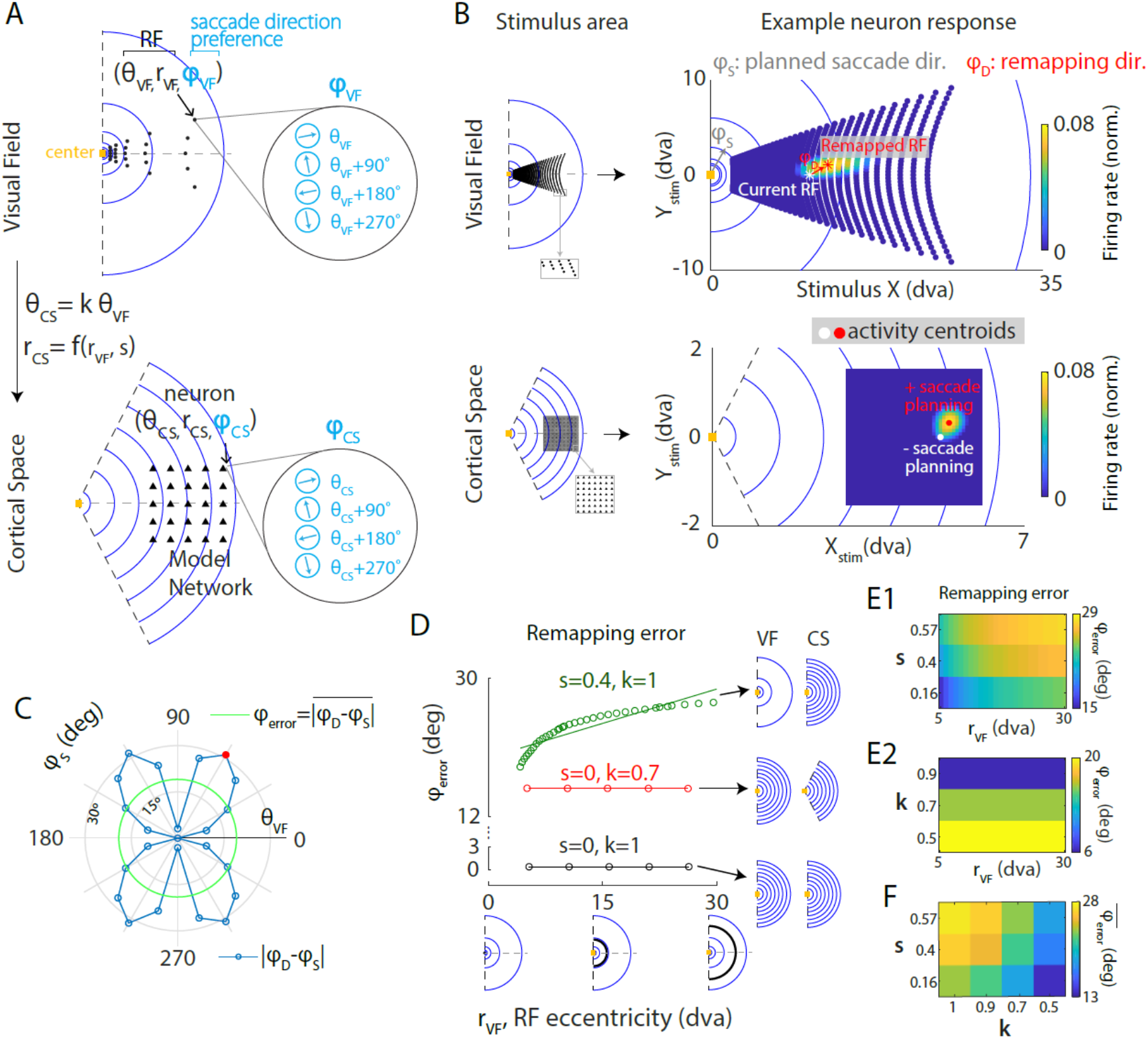
Network model of RF remapping under non-uniform cortical transformation. (**A**) Same as Fig. 1A1 but under a non-uniform cortical transformation. The RF location of each neuron (black dot, top row) is mapped to cortical space (black dot, bottom row) according to the implemented cortical transformation, which may involve: (a) eccentricity-dependent scaling of the RF radial coordinate (r_VF_), parameterized by s (see Methods), and (b) contraction of the polar angle (θ_VF_) with extent k. (**B**) Neural response with and without saccade planning. Left: Stimulation area in visual and cortical space. Visual stimuli were presented at different locations in the grid. Right: Steady-state response of neurons with identical spatial RFs (r_VF_, θ_VF_) during saccade planning, as a function of stimulus location. Dots/asterisks: population activity centroids and their corresponding coordinates in the visual field, with (red) and without (white) saccade planning. (**C**) Error in remapping direction for example neurons in (**B**) across various saccade directions (φ_S_), quantified as the difference between the direction of RF shift (φ_D_) and the saccade direction. Red dot: planned saccade direction simulated in (**B**). (**D**) Average error in remapping direction (φ_error_) as a function of RF eccentricity for different cortical transformations characterized by combinations of s and k. Black: uniform transformation. Green: eccentricity-dependent cortical magnification (CM) only. Red: polar angle contraction only. Circles: numerical simulations. Lines: linear fits (fitted slopes: 0.35 for s = 0.4, k = 1; 0 for other parameter combinations). (**E**) Dependence of remapping direction error on RF eccentricity. **E1**: CM only fs >0, k = 1 ). **E2**: Polar angle contraction only ( k < 1, s = 0 ). (**F**) Error of remapping direction averaged across RF eccentricities 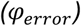 as a function of s and k. In (**E-F**), results were generated from fitted functions based on numerical results described in (**D**) (see Methods).

Similar to the model with uniform cortical transformation, we estimated the current RFs (CRF) and remapped RFs for the network (Fig. 4B, right; Methods). Given the rotational symmetry of the implemented transformation, we only simulated square grids of neurons with fixed network sizes on the cortical sheet that were symmetric about the 0° polar axis (Fig. 4B, left). By simulating the network at different cortical locations, we investigated the remapping performance of neurons with varying RF eccentricities.

### Non-uniform cortical transformation introduces remapping errors

We first examined whether the non-uniformity of cortical transformation introduced systematic errors in remapping direction. Saccade planning induced RF shifts in a direction (*φ*_*D*_, Fig. 4B) that deviated from the intended saccade direction (*φ*_*S*_; Fig. 4B), indicating a mismatch in remapping direction. The remapping error, defined as the angular difference between the direction of the RF shift (*φ*_*D*_) and the direction of planned saccade (*φ*_*S*_), depended on the planned saccade direction (Fig. 4C). Notably, it vanished when the saccade direction aligned with the RF polar angle (*θ*_*VF*_; Fig. 4C). This result is consistent with the intuition that under such a scenario, the axis of CD-induced imbalance in neural activity in cortical space corresponded to the axis in the visual field that aligned with the RF polar angle. Consequently, neuronal responses to visual stimuli symmetric about this axis remain identical, yielding remapped RFs along the axis and zero directional error. Furthermore, this result suggests that for a given planned saccade, neurons with RF polar angles aligning with the planned saccade direction (*θ*_*VF*_ = *φ*_*S*_) exhibit optimal remapping performance (*φ*_*D*_ = *φ*_*S*_).

### Both eccentricity-dependent cortical magnification and polar angle contraction, when present alone, increase remapping errors

We next investigated the role of CM and polar angle contraction separately in introducing remapping error. We first varied CM without polar angle contraction ( *k* = 1 ) and characterized remapping performance by estimating mean angular remapping error (*φ*_*error*_ ). Compared to uniform cortical transformation, CM (*s* > 0) resulted in larger remapping errors (Fig. 4D). The errors obtained through model simulation were well fitted by generalized power functions (Fig. S3B; Methods), allowing us to estimate remapping error for neurons with arbitrary CRF eccentricities. Moreover, the error magnitude increased with both CRF eccentricities of remapping neurons (*r*_*VF*_ ) and the RF size-eccentricity slope (*s*) (Fig. 4E1). When we varied the polar angle contraction alone (*k* < 1, *s* = 0), it resulted in remapping errors that were invariant to the CRF eccentricities of remapping neurons and increased with the extent of contraction, *k* (Figs. 4D, 4E2, and S3A).

### Polar angle contraction rescues remapping errors in the presence of eccentricity-dependent cortical magnification

We next sought to understand how CM and polar angle contraction interact to impact the quality of remapping. For each combination of parameters characterizing these two geometric transformations ( *s* and *k* ), we quantified the corresponding remapping performance by averaging remapping errors across both saccade directions and CRF eccentricities (Methods). In contrast to the exacerbating effect of polar angle contraction alone (*k* < 1, *s* = 0) on remapping errors (Figs. 4E2 and S3A), contraction reduced remapping errors when combined with CM (Fig. 4F) at all eccentricities (Fig. S3B). This result suggests polar angle contraction as a compensatory mechanism that mitigates the negative impact of non-uniform CM on remapping quality.

### Remapping errors increase with CRF eccentricity in V2

We again turned to the experimental data from area V2 (Fig. 2A-C) to test the model prediction that remapping direction errors should increase with CRF eccentricity. For each neuron, we characterized the stable remapped RFs and the corresponding RF shifts during saccade planning (Fig. 2I; Methods). Consistent with the model prediction, remapping direction errors increased with CRF eccentricity in both monkeys (Fig. 5), thus providing further support for the proposed theoretical framework. Because we previously showed that PS strength decreased with CRF eccentricity (Fig. 2H) and reduced remapping errors (Fig. 2J), we sought to examine whether the identified effect of CRF eccentricity on remapping error was confounded by PS strength. For neurons included in the error analysis (Figs. 2J and 5) PS strength and CRF eccentricity were negatively correlated (Fig. 2H). However, a residual-based regression analysis that regressed PS strength out from CRF eccentricity and remapping error and then regressed the resulting residual against each other yielded a significant effect (monkey M: N=55, r=0.29, p=0.03; monkey D: N=31, r=0.37, p=0.04), suggesting that the effect of CRF eccentricity on remapping accuracy is independent of variations in PS strength.

**Figure 5.**
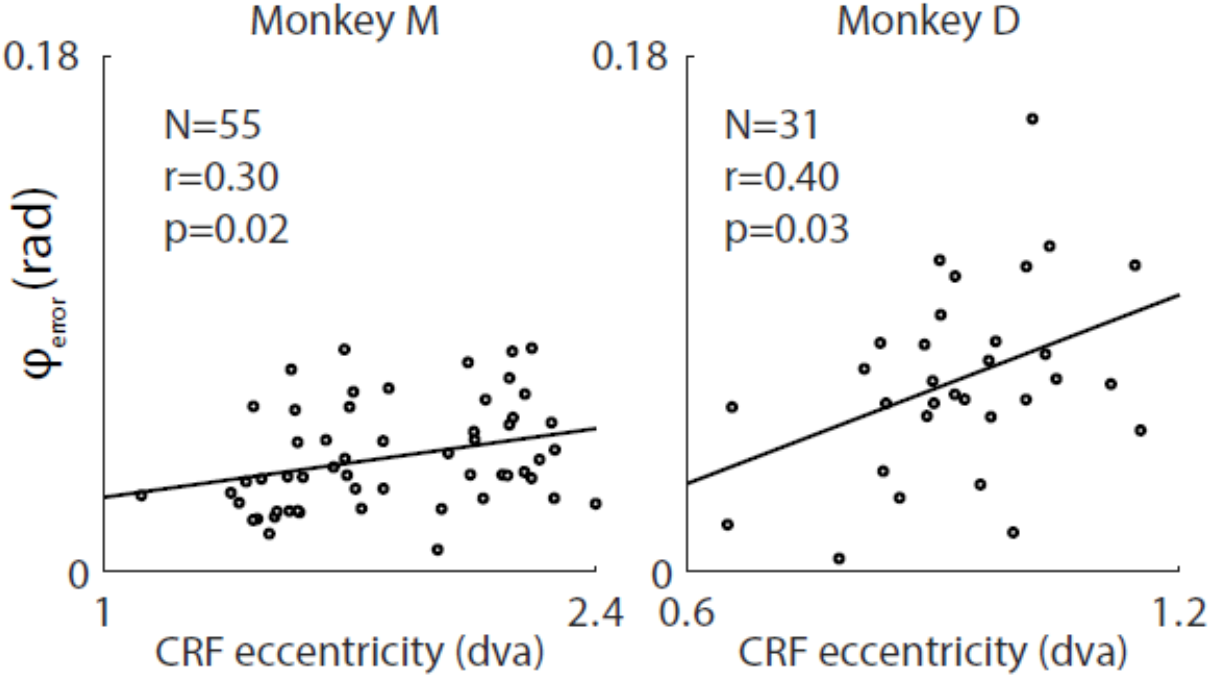
Relationship between CRF eccentricity and the error of remapping direction in V2. Remapping direction error, φ_error_ (defined in Fig. 2I), averaged across saccade targets as a function of CRF eccentricity for each monkey (see Methods). Black line: least squares fit to the data (slope: 0.05 for monkey M, 0.32 for monkey D).

### Geometric distortions underlie remapping errors in non-uniform cortical transformations

The dependence of remapping errors on CRF eccentricities, along with the competing influence of CM and polar angle contraction on remapping quality, led us to formulate the following hypothesis: remapping errors arise from geometric distortions introduced by non-uniform transformation from the visual field to cortical space, with greater distortions resulting in larger errors. To test this hypothesis, we quantified the distortion between the visual field and cortical space using the normalized Procrustes Distance (Fig. 6A, Methods), for various parameter combinations (*s* and *k*) characterizing the extents of non-uniform cortical transformations explored in the model. This distance measure characterizes the minimal difference between two shapes – each defined by a distinct set of points – by identifying the optimal mapping between them under translation, rotation, and scaling transformations.

**Figure 6.**
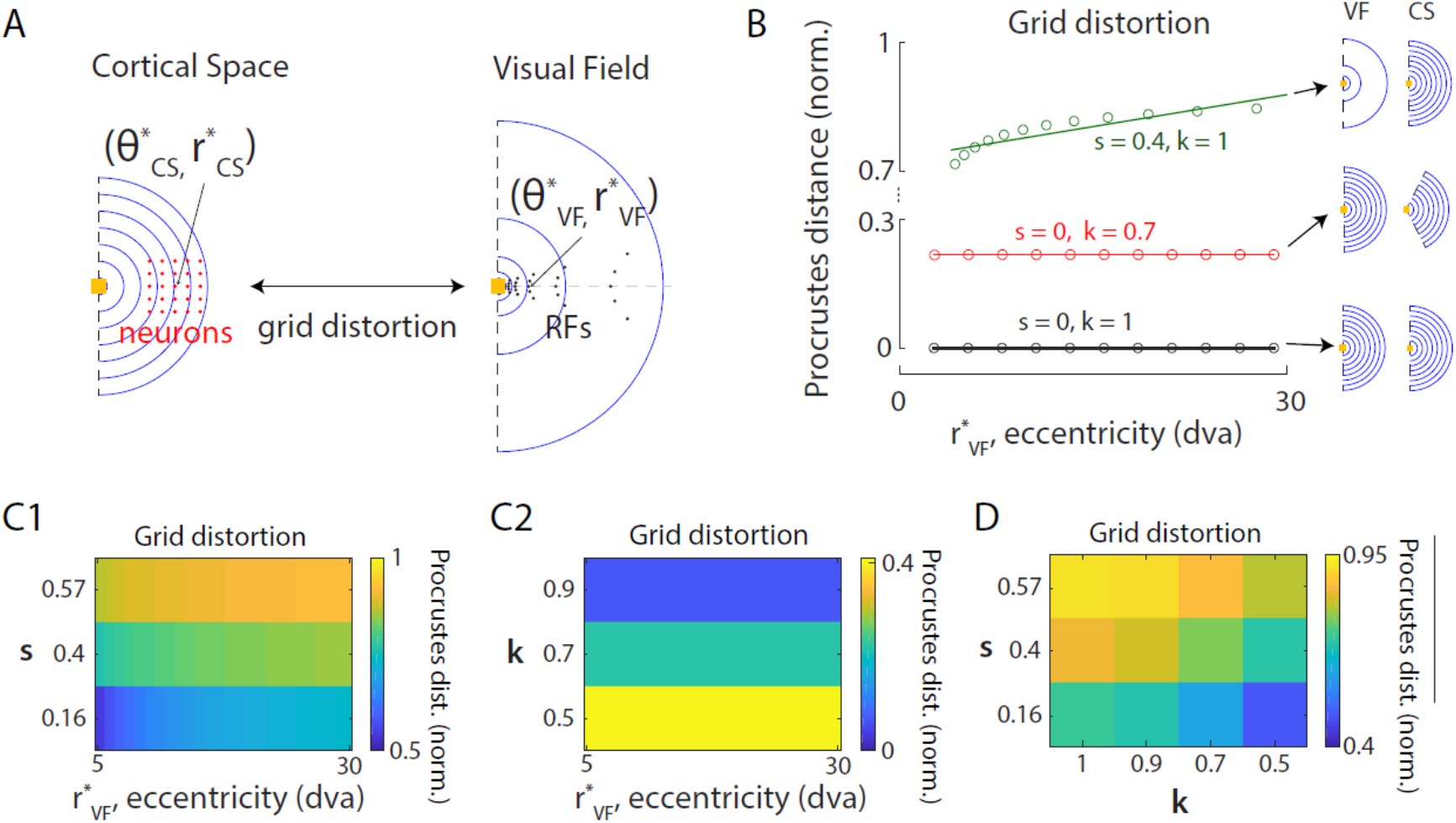
Representation distortion between the visual field and cortical space due to non-uniform transformation. (**A**) Illustration of distortion analysis. Distortion is computed by comparing the geometric arrangement of network neurons (red dots) in cortical space to the arrangement of their RF locations (black dots) in the visual field following a given cortical transformation. The network location is defined by the RF location of its center neuron, with polar coordinates as 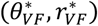 . (**B**) Grid distortion quantified by normalized Procrustes Distance (see Methods) as a function of network eccentricity 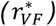 for different cortical transformations characterized by combinations of RF size-eccentricity slope (s) and the extent of polar angle contraction (k). (**C**) Grid distortion as a function of network eccentricity. **C1**: CM only (s > 0). **C2**: Polar angle contraction only (k < 1). (**D**) Grid distortion, averaged across network location eccentricities, as a function of s and k. In (**C-D**), the results were generated from the fitted functions based on numerical results described in (**B**) (see Methods).

We examined square grids of neurons with fixed network sizes symmetric about 0° polar axis, and quantified the distortion between their cortical locations and the corresponding RF locations in the visual field (Fig. 6A). As expected, uniform cortical transformation (*s* = 0 and *k* = 1) resulted in a zero normalized Procrustes Distance with the original shape, since every point in the visual field is uniformly scaled when mapped to cortical space (Fig. 6B). In contrast, CM induced distortion whose magnitude as a function of the eccentricity of network location 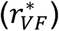 was well fitted by a generalized power function (Fig. 6B; Methods). Moreover, the magnitude of distortion increased with both eccentricity and the extent of cortical distance-dependence of CM, governed by *s* (Fig. 6B, C1). Thus, the effects of CM on remapping errors, as well as the dependence on eccentricity, are qualitatively replicated by the distortion analysis.

Additionally, polar angle contraction reduced distortion magnitude in the presence of CM (Figs. 6D and S4B), although the magnitude of distortion due to polar angle contraction alone was invariant to eccentricity and increased with the extent of contraction (Figs. 6C2 and S4A). This result mirrors the identified interplay between CM and polar angle contraction in affecting remapping quality. Taken together, these findings indicate that the remapping errors arising from non-uniform cortical transformation can be explained by the nature of distortion from the visual field to cortical representation. Eccentricity-dependent logarithmic CM introduces distortion along radial directions, with larger amplitudes at higher eccentricities. Such distortion can be counteracted by the distortion along tangential directions introduced through polar angle contraction, thereby rescuing remapping quality. This interplay demonstrates how biological circuits can balance the competing demands of foveal-biased CM and predictive remapping accuracy within the constraints of finite cortical space.

## DISCUSSION

In the present study, we establish a generalized recurrent 2D network architecture that explains how the primate visual system can achieve stable and coherent predictive remapping in the presence of non-uniform selectivity, noisy inputs and non-uniform distortions in cortical representation. By combining computational modeling with dense electrophysiological recordings in macaque area V2, we demonstrate that the integration of pre-saccadic corollary discharge signals by conjunctively tuned neurons is sufficient to maintain spatial constancy across saccades. More specifically, we show that a CD signal with two competing components – a selective excitation to boost neurons tuned to the planned saccade direction, and a non-selective suppression to brake runaway recurrent excitation – ensures stability and fidelity of predictive RF remapping, while recurrent connectivity maintains local receptive field relationships against sensory and motor noise. Finally, our findings reveal a novel role for countervailing cortical distortions in systematically mitigating eccentricity-dependent remapping errors. Simultaneous population recordings in V2 from macaques performing a cued saccade task, validated not only the model predictions on saccade direction selectivity, but also the factors that can influence remapping errors. Together, these results provide a unified circuit mechanism showing how the brain fundamentally reconciles the competing demands of high-acuity foveal vision and seamless perceptual continuity during saccades.

### Dual role for pre-saccadic suppression in remapping

Pre-saccadic suppression of neuronal activity has been consistently observed across visual areas. While this phenomenon is well recognized for modulating neuronal sensitivity to blunt the perception of saccade-induced motion and thereby preventing misinterpretation of rapid image shifts (4), our model predicts expanded functional roles. Modeled as a suppressive component of corollary discharge signal, PS further enhances visual stability by facilitating stable RF remapping, which predictively aligns the pre- and post-saccadic spatial representations. Specifically, it counteracts the runaway excitation effects arising from a key feature of our model – saccade direction selectivity – to maintain stable and robust remapping. Additionally, modeled as an untuned signal, PS helps remapping accuracy across visual space by sharpening the saccade direction selectivity of local neuronal population throughout cortex (Fig. 1B1 & S1D1). Similar to the matching time courses of perceptual suppression and neural activity suggesting a role of saccadic suppression in dampening perception (23), the overlapping onset of pre-saccadic neuronal suppression and RF remapping in V2 – both beginning approximately 40 ms before saccade onset (9) – offers support for our prediction. Whether similar temporal coupling occurs in other visual areas remains an open question for future investigation. Moreover, our prediction offers an interpretation for the observed correlation between pre-saccadic suppression of neural activity and the RF remapping phenomenon across many visual areas, as both phenomena are weak in V1 (4, 8, 19, 26) but robust in higher visual areas such as V2 (8, 9) and V4 (15, 21). Notably, our prediction that suppression facilitates remapping does not imply it is a necessary condition; some areas may operate in a less excitable regime and achieve remapping even in the absence of suppression (49).

### A low-dimensional manifold of population RFs during remapping

The conserved cell-cell RF relationships we identified from cortical recordings during remapping can be interpreted through the lens of a low-dimensional manifold. Specifically, despite the high-dimensional ambient space of population RF coordinates (2*N* dimensions for *N* neurons with RFs in 2-D space), an internal structure confines the population RFs to a 2-D retinotopic manifold during saccade planning, consistent with our model prediction. This conserved cell-cell relationship is unlikely to be inherited from upstream area V1, where the proportion of neurons exhibiting predictive remapping is significantly lower (8). Furthermore, since the transient deformation of cell-cell RF relationships occurs only along the saccade vector (Fig. 3E, F), this can be interpreted as saccade planning temporarily displacing the network state off the 2-D manifold of RF states (defined by retinotopy), specifically along the planned saccade direction. Due to the attractive property of the manifold, the network state eventually returns to the manifold at a new location, with each neuron’s RF shifted by a similar distance in the planned saccade direction.

### Relationship to other models of dynamic allocentric representations

In a broader context, the computation underlying predictive RF remapping can be understood as the updating of allocentric spatial representations to accommodate shifts in the reference frame, guided by egocentric signals encoding these shifts. Our model incorporates two ingredients proposed in prior modeling work exploring analogous mechanisms, primarily in the context of path-integration in head-direction cells and grid cells of the medial temporal lobe (50-54). First, neurons are conjunctively selective to both spatial properties (e.g., visual field location in remapping, spatial location/direction in path-integration) and egocentric shifts in the reference frame (e.g., impending saccades in remapping, velocity in path-integration; Figs. S1A and 1B1). Second, the network includes an asymmetric recurrent connectivity pattern, where the profile of asymmetry depends on the presynaptic neuron’s selectivity to egocentric shifts (Fig. S1A). The latter assumption has been validated by a recent study of grid cells in the medial entorhinal cortex (55) and remains to be examined in visual areas.

Our model differs from classical path-integration frameworks in two ways. First, whereas standard path-integration models assume neurons are arranged topographically by spatial preference, our model incorporates non-uniform cortical transformations, which disassociates neural distance and RF similarity. Second, we model visual areas – consistent with prior evidence (44, 56) – as operating in a dynamical regime of amplification, characterized by silence in the absence of feedforward inputs (visual stimuli here) and a prolonged response following the disappearance of an input. This contrasts with the continuous attractor regime adopted in previous path-integration models, which is marked by self-persisting activity. Our focus on stationary states under tonic external inputs allowed us to omit explicit excitatory-inhibitory (E-I) separation (44).

### 2-D network architecture as a basis for investigating remapping quality

While prior models have captured the fundamental mechanistic aspects of predictive remapping primarily in 1-D networks with CD-gated directional connections (21, 22), evaluating remapping quality across a continuous visual space requires investigating these circuit mechanisms in a 2-D cortical patch. In addition to enabling precise characterization, a 2-D framework allows for the incorporation of non-uniform cortical transformation – a feature grounded in empirical findings across visual areas. This modification introduces a dissociation between neural separation (i.e., cortical distance) and RF difference, such that synaptic connectivity no longer depends uniformly on RF similarity. Our study reveals that non-uniform cortical transformation introduces systematic remapping errors, rooted in the geometric distortion between visual space and its cortical representation.

### Potential extensions

Although the present model restricts noise to the network inputs— which can be interpreted as variability in either visual stimuli or corollary discharge signals—noise manifests in different ways in the brain, such as noise in intrinsic neuronal properties. Investigating these alternative noise sources within our established 2-D framework remains an open direction for future research.

In this study, our modeling and empirical analyses focused on forward remapping along the saccade vector rather than convergent remapping toward the saccade target. Area V2 serves as an ideal area to test the predictions of this model, as a previous study from our laboratory utilizing the same dataset demonstrated that a clear majority (73%) of units exhibit robust forward remapping, with no evidence of widespread convergent remapping (9). This allowed us to empirically validate the network’s core mechanisms in a visual area where forward remapping is naturally isolated.

Building upon this baseline, a future direction is to extend the framework to accommodate convergent remapping, which can be achieved by incorporating an attention-modulated input, as explored in a recent paper (30). That study also proposed that the net input to the network could be modeled as a linear combination of corollary discharge inputs driving forward shifts and attentional modulation inputs driving convergence. Thus, while our model captures the primary operational principles tested here in area V2, its architecture provides a scalable foundation that can be readily extended to investigate how the interactions of dual-shift mechanisms affect remapping quality in future exploration.

### Implications for experiments

As we show, during saccade planning, the impact of CD signals on network dynamics, and the consequent remapping accuracy, depends on both the saccade direction (Fig. 4C) and the network location as indexed by RF eccentricity (Figs. 4D-E and 5). The directional dependence is consistent with prior findings in area V4, where saccades directed toward the RFs of recorded neurons elicit stronger direction-specific traveling waves, typically propagating from the fovea to the periphery (57). Thus, comprehensive empirical investigations should systematically assess remapping accuracies for neurons with varying RF locations and evaluate perceptual changes for stimuli at varying eccentricities, with both analyses conducted under variations in saccade direction.

By identifying the dependence of remapping errors on the two non-uniform geometric characteristics – eccentricity-dependent CM and polar angle contraction – our study establishes a framework to predict remapping accuracy across visual areas with varying extents of these characteristics. Across visual areas along the ventral stream, the CM factor (CMF) is well characterized by an inverse function of eccentricity (34, 39, 58-60), with its integral reflecting logarithmic CM. The scale of this function progressively decreases from V1 to V2 to V4 (34, 37-39), resulting in a stronger cortical distance-dependence of CM, marked by increasingly rapid decay rates of CMF. While this heightened dependence alone is predicted by our model to induce greater remapping errors along the ventral stream, a comprehensive prediction should also account for the extent of polar angle contraction, which remains to be further characterized experimentally. Our model suggests that the extent of contraction along the ventral stream may increase to compensate for the exacerbating effect of increasing distance-dependence of CM on remapping accuracy.

## ACKNOWLEDGEMENTS

This research was supported by NIH/NEI R01 EY034605 and R00 EY025026 to MPJ, NIH/NEI R01 EY032555, NARSAD Young Investigator Grant, Ziegler Foundation Grant and Yale Orthwein Scholar Funds to ASN, Kavli Postdoctoral Fellowship and Swartz Postdoctoral Fellowship in Theoretical Neuroscience to XX, NIH/NINDS training grants T32-NS007224 and T32-NS041228 to SD, and by an NIH/NEI core grant for vision research P30 EY026878 to Yale University. We want to thank the veterinary and husbandry staff at Yale for excellent animal care.

## AUTHOR CONTRIBUTIONS

XX & MPJ conceptualized the project. XX conducted the model simulations and analysis. SD & ASN designed the experiments, and ASN supervised the data collection. SD collected data and performed preliminary analyses. XX conducted the physiological analysis. MPJ supervised computational modeling and data analysis. XX, ASN & MPJ wrote the manuscript.

## DATA AVAILABILITY

All data reported in this paper will be shared upon publication.

## MATERIALS & METHODS

### Network model with uniform cortical transformation

The activity of neuron *i* was characterized by rate-based dynamics:

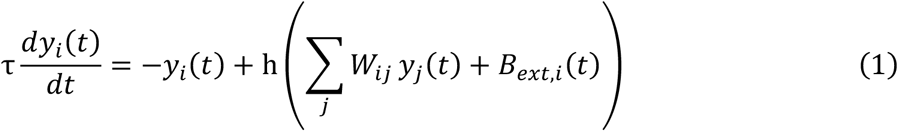

where τ was the time constant of neural response, *W*_*ij*_ the synaptic weight from neuron *j* to neuron *i, B*_*ext,i*_(*t*) the external input, and *h* the neural transfer function taken as a simple rectification nonlinearity: h*(x*) = *ax* for *x* > 0 and was 0 otherwise.

We assumed neurons with receptive fields (RF) 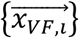 in the visual field were uniformly arranged at *N*^2^grid locations on a 2-D cortical sheet. Neuron *i* was located at 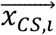, The network center was located at 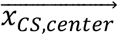. 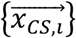 formed a uniformly spaced square grid spanning from 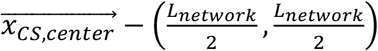 to 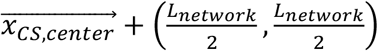 . The relationship between 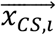 and 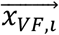 was defined by 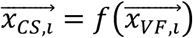. *f* defined the transformation from the visual field to cortical space:

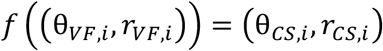

with

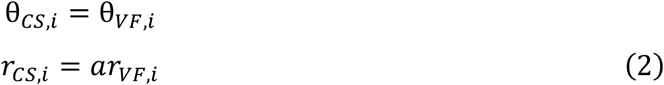

where (*θ*_*VF,i*_, *r*_*VF,i*_) was the polar coordinate of 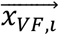 in the visual field, (*θ*_*CS,i*_, *r*_*CS,i*_) the polar coordinate of 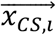 on the cortical sheet with the origin representing the center of visual space. Since this transformation uniformly scaled visual space, the topology of visual space was preserved on the cortical sheet.

At each location, there were 4 neurons with the same RFs and distinct saccade direction preferences (SDP) {φ_*VF,i*_}. Therefore, there were 4*N*^2^neurons in the network. The SDPs of the neurons at location 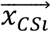 were restricted to 0°, 90°, 180°, 2−0°. While the SDPs were restricted to four directions for the convenience in modeling, in the visual cortex, the SDPs might span the continuum 0,2π (46). The SDP of a neuron was used to determine (1) the direction in which its outgoing weights were shifted and (2) the corollary discharge input it received.

The synaptic weight between a pair of neurons depended on their spatial distance and the SDP of the presynaptic neuron:

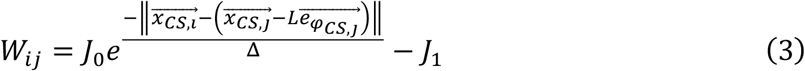

where 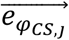 was the unit vector pointing along *φ*_*CS,j*_. The outgoing weight matrix of neuron *j* had a center-surround shape centered at the shifted location 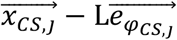, where *φ*_*CS,j*_ = *g*(*φ*_*VF,j*_ ) was the equivalent SDP on the cortical sheet:

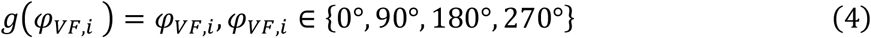

The external input consisted of three components:

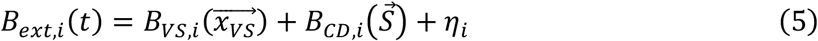

The input due to a visual stimulus presented at location 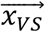 was

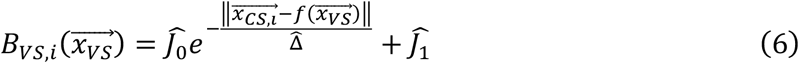

In the parameter regime of stability, this input alone induced an activity bump centered at the corresponding location of visual stimulus on the cortical sheet.

The corollary discharge (CD) input during the planning of a saccade *S* was

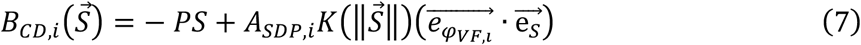

where *PS* was the level of pre-saccadic suppression, 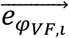 the unit vector pointing along *φ*_*VF,i*_, 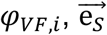 the unit vector pointing along the saccade direction (*φ*_*S*_). *A*_*SDP,i*_ was the strength of saccade direction bias in *φ*_*VF,i*_, which could be uniform or vary with *φ*_*VF,i*_ to mimic the non-uniform distribution of SDP, as explored in Figs. 1B4 and S1D. The strength of CD input also depended on the magnitude of saccade 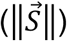 through *K*. In this paper, we focused on the accuracy of remapping direction and therefore took 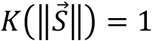 in all simulations. One may match a value of *K* to any magnitude of saccade (31).

In addition, a spatially varying noise term, η_*i*_, was introduced into the inputs (Fig.1B3). This term was independent across neurons and drawn from a Gaussian distribution with zero mean and standard deviation *σ*_*noise*_. This way of implementing noise can be interpreted either as noise within the visual stimulus or as noise within the CD signal. Note that this noise term is absent in most figures to focus on the underlying network mechanism of remapping and isolate the effect of other elements, except for Fig.1D. In the model, *L, A*_*SDP,i*_ and 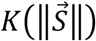 allowed saccade direction to couple to the network dynamics, characterized by the shift of a bump of neural activity on the cortical sheet compared to the scenario without saccade planning: when 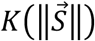 were non-zero, neurons with identical RFs but distinct SDPs received different strengths of CD inputs 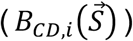, thus generating an imbalance at locations with non-trivial neural activities. When *L* was non-zero, this imbalance caused the population activity to propagate in the opposite direction of the planned saccade 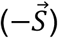. Together with visual stimulus input 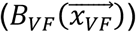, which tended to fix the activity bump at the cortical location representing the location of the visual stimulus 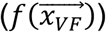, a stable bump could be obtained in the direction opposite to the saccade relative to 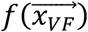. The stability required the shift *L* to be small. In this paper, we set L to be twice the distance between adjacent neurons. The magnitudes of *L, A*_*SDP,i*_ and 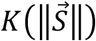 multiplicatively determined how far CD inputs shifted the bump from the location representing visual stimulus, and thus controlled the magnitude of remapping.

### Modifications for network model with non-uniform cortical transformation

Parameters for the neuron and network structure were kept fixed as in the scenario with uniform transformation (see Table 1).

**Table 1.**
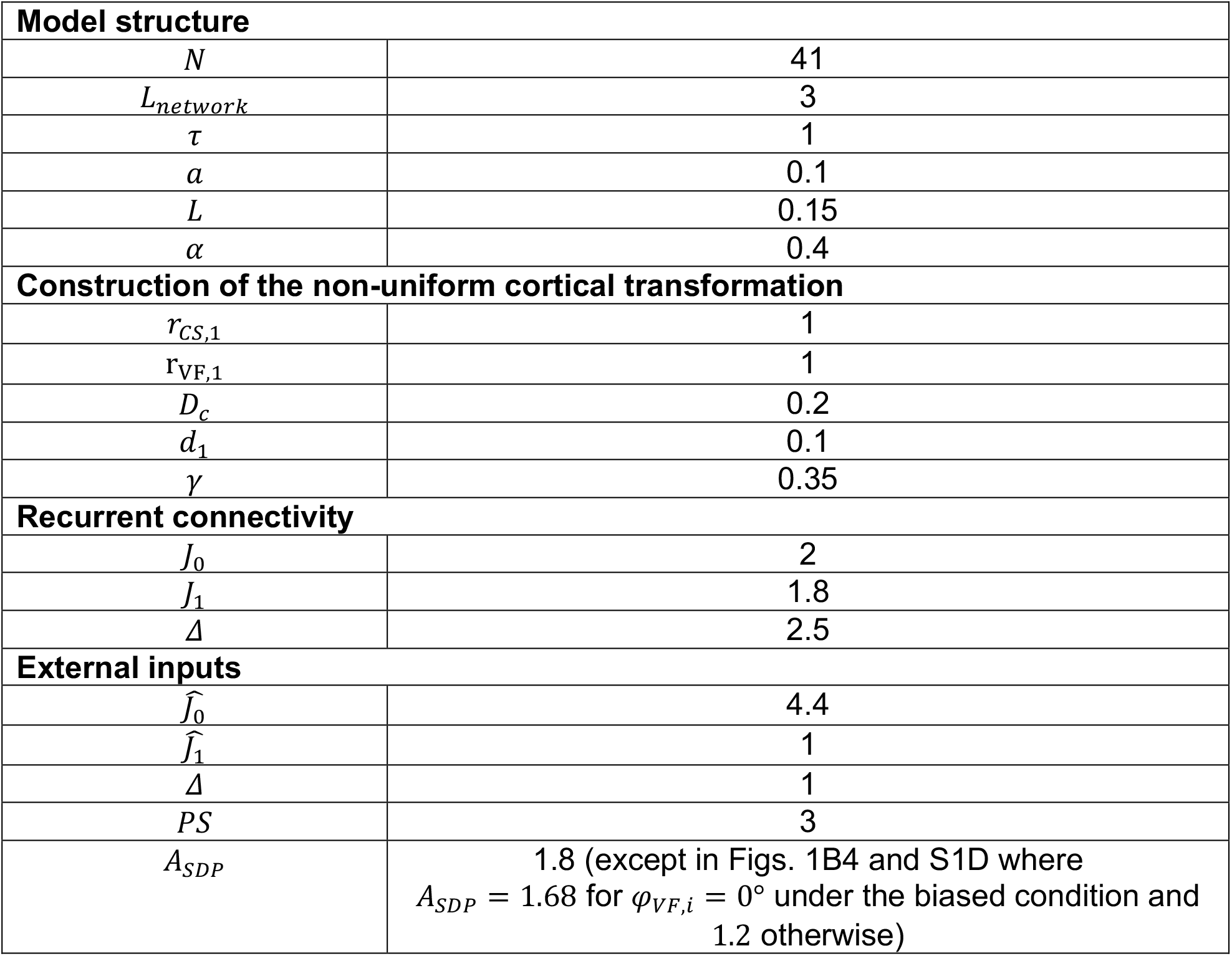
Parameters in the network model.

| Model structure |  |
| --- | --- |
| $N$ | 41 |
| $L_{network}$ | 3 |
| $\tau$ | 1 |
| $a$ | 0.1 |
| $L$ | 0.15 |
| $\alpha$ | 0.4 |
| Construction of the non-uniform cortical transformation |  |
| $r_{CS,1}$ | 1 |
| $r_{VF,1}$ | 1 |
| $D_c$ | 0.2 |
| $d_1$ | 0.1 |
| $\gamma$ | 0.35 |
| Recurrent connectivity |  |
| $J_0$ | 2 |
| $J_1$ | 1.8 |
| $\Delta$ | 2.5 |
| External inputs |  |
| $\hat{J}_0$ | 4.4 |
| $\hat{J}_1$ | 1 |
| $\Delta$ | 1 |
| $PS$ | 3 |
| $A_{SDP}$ | 1.8 (except in Figs. 1B4 and S1D where $A_{SDP} = 1.68$ for $\varphi_{VF,i} = 0^\circ$ under the biased condition and 1.2 otherwise) |

#### • Cortical transformation (*f*)

For the network with non-uniform cortical transformation, we implemented two geometric characteristics of retinotopic maps observed across visual areas. First, the space of polar angles in visual space was represented with a shrinkage in the cortex. Second, RF size increased linearly with eccentricity, with the slope increasing from V1 to V2 to V4 (37, 38).

The implementation of the linear RF size-eccentricity relationship captured the logarithmic cortical magnification (CM), as shown in (48). Briefly, we considered a line of uniformly spaced neurons on a cortical sheet with RFs along an iso-polar contour. The radial coordinate of the *n*th neuron, *r*_*CS*,n_, was given by

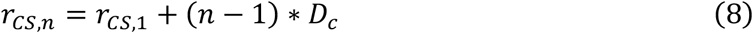

where *D*_c_ was the distance between adjacent neurons and *r*_*CS*,1_ the radial coordinate of the first neuron. The eccentricity and diameter of the RF of the *n*th neuron were r_*VF*,n_ and *d*_n_, respectively. The RF of the first neuron was located at r_FG,1_ = 1 with a diameter *d*_1_ = 0.1. Let *s* be the slope of the linear function relating RF radii to eccentricity *r*_*VF*_, which was taken as 0.16, 0.4, 0.57 to model the retinotopy maps of V1, V2 and V4, respectively (37, 38) (Fig. S5A):

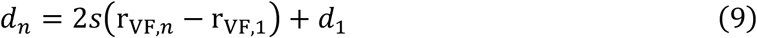

Here the slope *s* was assumed to be positive since a zero value of *s* would result in uniform cortical transformation as characterized by Eq. 2.

Let *γ* be the proportion of RF diameter overlap between adjacent neurons, which satisfied 0 < *γ* < 1 and was taken as 0.35 in our model:

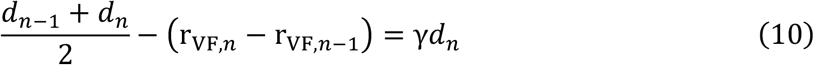

Implicit in Eq. 9 and 10 were r_FG,n_, *d*_n_ as functions of r_FG,*n*−1_, *d*_*n*−1_. The RF eccentricity r_FG,*n*_ of the *n*th neuron, was given by:

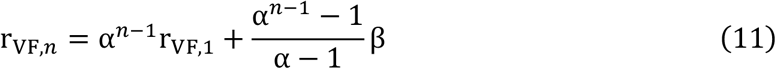

where 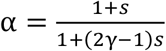 and 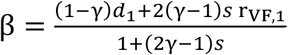. Since α > 1 for 0 < *γ* < 1, away from the immediate vicinity of r_FG,1_, *log*(r_FG,*n*_) could be written in an approximate form as

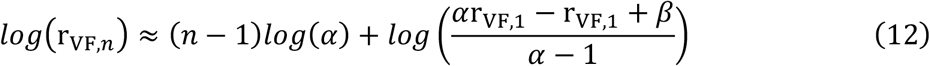

yielding an expression of *n* ™ 1 in terms of r_FG,*n*_. Plugging this into Eq. 8 gave r_CS,*n*_ as a logarithmic function of RF eccentricity r_*VF,n*_ (Fig. S5B):

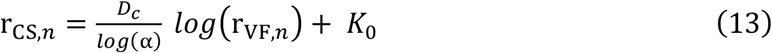

where

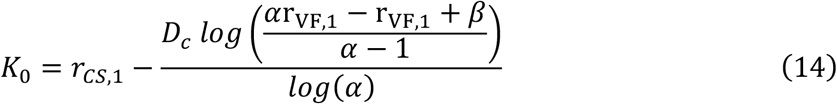

Together with the shrinkage of the RF polar angle, visual space was therefore represented on the cortical sheet by:

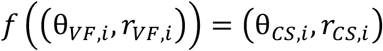

with

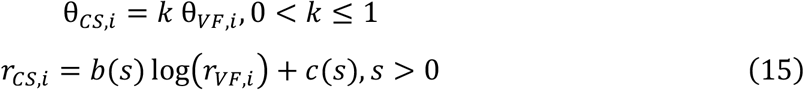

where *k* was the extent of the shrinkage of the angle space in visual space. 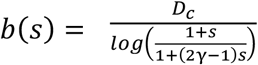 and *c(s*) was given by *K*_0_ (Eq. 14). Thus the CM factor (CMF), quantifying how the amount of cortical surface area dedicated to visual processing varies with the corresponding RF location in the visual field, was 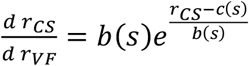. Since *b(s*) decreased with *s*, larger values of *s* corresponded to stronger cortical distance-dependence of CMF, marked by faster decay rates with *r*_*CS*_ (Fig. S5C-D). Due to the rotation-invariance of the implemented transformation, we only simulated square grids of neurons with fixed network sizes on the cortical sheet that were symmetric about the 0° polar axis.

#### Saccade direction preferences (SDP;{ *φ*_*VF,i*_})

The SDPs of neurons ( *φ*_*VF,i*_) at location 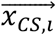 were restricted to {*θ*_*VF,i*_ + 0°, *θ*_*VF,i*_ + 90°, *θ*_*VF,i*_ + 180°, *θ*_*VF,i*_ + 270°}, which corresponded to directions along the radial axis (defined by the axis connecting the fixation point and the RF), and its orthogonal axis in visual space (Fig. 4A). The equivalent SDP on the cortical sheet was *φ*_*CS, j*_, where

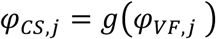

with

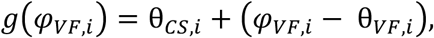

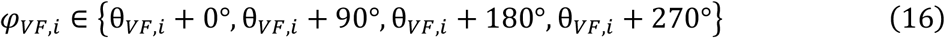

We reasoned that under the applied cortical transformation, the direction of an infinitesimal shift along the radial direction (*θ*_*VF,i*_) in the visual field was equivalent to a shift in radial direction (*θ*_*CS,i*_) in cortical space. This equivalence extends to the other three SDPs such that *θ*_*VF,i*_ + 90° → *θ*_*CS,i*_ + 90°, *θ*_*VF,i*_ + 180° → *θ*_*CS,i*_ + 180°, *θ*_*VF,i*_ + 270° → *θ*_*CS,i*_ + 270°.

### Simulation protocol

Periodic boundary conditions were applied to both networks with uniform and non-uniform cortical transformations. All simulations began with random initial conditions. The time step for numerical integration was set to *dt* = 0.1τ. The network was simulated for a sufficiently long duration (*T* = 1800*dt*) to reach its steady state, characterized by an activity bump. For the model incorporating noise into the network inputs (eqn(5)), 500 independent noise realizations were simulated to steady state to gather statistics.

The RF of a neuron was tested by presenting visual stimuli at the RF locations of each neuron to the network and recording the induced stable population activity under two conditions: (1) without saccade planning (*B*_*CD,i*_ = 0), (2) during the planning of a saccade in a specific direction. This process generated for each neuron a map of response as a function of the cortical representation of the visual stimulus location. The maps of neurons with identical cortical locations (hence identical RFs) were averaged, where the centroid was determined and mapped to visual space as the current RF under test condition (1), which was 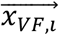 by definition, or the remapped RF under test condition (2) 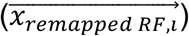. The RF shifts (*D*) of neurons representing 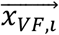 were calculated as 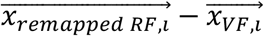. The direction of *D*, denoted as *φ*_*D*_, was referred to as the ‘remapping direction’. The error of remapping direction was calculated as the absolute difference between the remapping direction and the saccade direction (*φ*_*error*_ = |φ*error* = _*D*_ ™ *φ*_*S*_|). The cell-cell RF relationships were quantified by the mean relative pairwise difference in RF shifts: 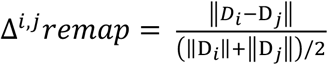 (Fig. 1C4). In Figs. 1A4 and 1C3, we computed this metric using neurons at nine cortical sites in a square grid with coordinates *x* ∈ {9.55,9.−,9.85} and *y* ∈ {™0.45, ™0.3, ™0.15} across various levels of recurrent connectivity strength ( Δ = 1.5,2.5,3.5). Under non-uniform cortical transformation (Fig. 4), we simulated square grids of neurons with fixed network sizes on the cortical sheet that were symmetric about the 0° polar axis. Since the range of CRF eccentricity captured by one network increased with *s*, where larger values of *s* corresponded to stronger cortical distance-dependence of CMF, different numbers of network locations were explored to cover a comparable range of RF eccentricities: ten locations (network center 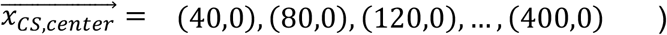) for *s* = 0, four 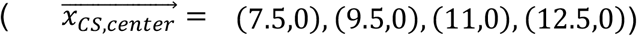 for *s* = 0.16, three 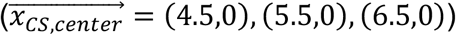 for *s* = 0.4, and one 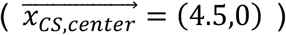 for *s* = 0.5− . Remapping performance, characterized by errors in remapping direction averaged across saccade directions, was determined for neurons with varying CRF eccentricities (Figs. 4D and S3). Due to the discretization of the network neurons and varying network locations explored, the captured values of CRF eccentricities were not aligned across *s* parameter values. To address this issue, the errors as functions of CRF eccentricities obtained through model simulation were fitted by generalized power functions: a linear relationship was fitted between the log of the derivative of errors 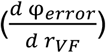 with respect to CRF eccentricity (*rVF* ), and the corresponding power function was the shifted to best match the numerical results, resulting in good fitting performance (see Fig. S3B for examples). This approach allowed us to estimate remapping errors for neurons with arbitrary CRF eccentricities (Fig. 4D-E). Furthermore, for each combination of *s* (governing the extent of cortical distance-dependence of CM) and *k* (extent of polar angle contraction), the corresponding remapping performance was characterized by remapping errors averaged across both saccade directions and a range of CRF eccentricities (from 5 dva to 55 dva) based on the fitted functions (Fig. 4F).

### Geometric distortion analysis

Under various combinations of the extents (*s, k*) of non-uniform cortical transformation incorporated into the model, we quantified the distortion of a shape when mapped from the visual field to cortical space using normalized Procrustes Distance (61). This distance characterizes the minimal difference between two shapes – each defined by a distinct set of points – by identifying the optimal mapping between them under translation, rotation, and scaling transformations.

Similar to the modeling study, we probed square grids of neurons located at 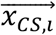 with fixed network sizes that were symmetric about 0° polar axis, computing the normalized Procrustes Distances between the shape defined by their cortical locations 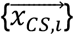 and the shape defined by the corresponding RF locations 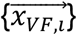 in the visual field (Fig. 6A). Specifically, the normalized Procrustes Distance was given by:

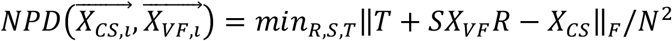

where *X*_*CS*_ denoted the matrix with each row being 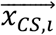 and similarly for *X*_*VF*_ ; *R, S, T* denoted the rotation, scaling and translation needed to best match the two shapes defined by 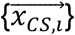 and 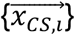, respectively; ‖∙‖_F_ was the Frobenius norm and *N*^2^the total number of neurons in the network. A greater *NPD* indicates larger distortion of cortical representation. Under uniform cortical transformation, 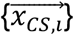 and 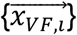 were completely overlapped under scaling and aligning, resulting in *NPD* 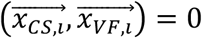,(Fig. 6B), implying no distortion.

We determined the magnitude of distortion across the 0° polar axis in both the visual field and cortical space. The magnitude of distortion as a function of eccentricity, given by the RF eccentricity of the neuron at the center of the network 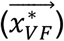, was fitted by a generalized power function, similar to the approach used for estimating the remapping error as a function of CRF eccentricities in the modeling study.

### Data Collection

## Surgical procedures

Low-profile titanium recording chambers were implanted in two rhesus macaques (62-64). Using preoperative structural MRI and sulcal reconstruction, the chambers were targeted over the lunate sulcus, allowing access to Area V2 (left hemisphere in monkey M, right hemisphere in monkey D). The native dura mater overlying this region was removed and replaced with a transparent silicone artificial dura (AD) (62-64). The AD allowed for visualization of area V2 and facilitated the targeting of electrode arrays. All procedures were approved by the Yale University Institutional Animal Care and Use Committee and conformed to NIH guidelines.

### Electrophysiology

Prior to a series of recordings, we electroplated (nanoZ, White Matter LLC) 64-channel electrode arrays (“laminar probes,” NeuroNexus Technologies, Inc., 2 shanks, 32 channels/shank, 70µm spacing between sites, 200µm between shanks) with PEDOT (poly(3,4-ethylene dioxythiophene)). At the beginning of each recording session, a laminar probe was lowered into Area V2 with an electronic micromanipulator (Narishige Inc.). Visual inspection of the cortical surface through a surgical microscope (Leica Microsystems) allowed for precise targeting of these probes, as well as continuous monitoring of electrode entry. To position the probe within the brain, we first penetrated the AD, arachnoid, and pia by moving the probe downward at a high speed (>100µm/s). After the tip of the probe entered the cortex, we inserted the remainder of the probe at a slow speed (2µm/s). Once the entire probe was in the brain, we slowly (2µm/s) relieved the pressure on the brain by retracting the probe upward, relieving pressure without moving the probe relative to the cortex.

Electrical signals from the laminar probe were collected at 30kHz, digitized on a 64-channel digital headstage, and sent to the recording system (RHD Recording System, Intan Technologies). Action potential waveforms were extracted offline using Kilosort2 (65, 66) with default settings (threshold = [10, 4], lambda = 10, AUC for splitting = 0.9) and manually sorted into single and multi-unit clusters (Phy; 65, 66). Units with a maximum waveform amplitude preceding the trough were classified as axonal spikes and excluded. Recordings were collected over the course of 17 sessions (8 in monkey M, 9 in monkey D). In total, 923 single units were recorded (461 in monkey M, 462 in monkey D). Only single units that were visually responsive, exhibited substantial response (>3 Hz) and had a significant spatial receptive field, as determined by a chi-squared test, were considered for subsequent analysis.

### Behavioral Control and Eye Tracking

We controlled behavioral experiments using NIMH MonkeyLogic (67). Eye position and pupil diameter were continuously sampled at 120Hz (ETL-200, ISCAN Inc.) and sent to the behavioral control system. Stimuli were presented on a monitor (BenQ XL2411; 60 Hz refresh rate) positioned 57cm from the monkey. Tolerance windows for fixation control were 1 degree of visual angle.

### Receptive field of the recorded column

We mapped RFs of the column by presenting Gabor patch stimuli (2-4 cycles/degree, 0.5-1.5 degree Gaussian half-width, 100% luminance contrast) on a square grid spanning the visual quadrant of interest (lower right in monkey M, lower left in monkey D) while the subject maintained fixation on the center of the screen. Grid spacing parameters were optimized for each session based on receptive field eccentricity and ranged from 0.25–1.0 degrees of visual angle (dva). A single Gabor was presented at one of six orientations (0, 30, 60, 90, 120, 150°) and at a grid location, both chosen at random, on each frame of stimulus presentation (60 Hz). Stimulus-evoked local field potential (LFP) power at each grid location on each recording channel was smoothed with a Gaussian kernel, and the peak location was defined as the RF center.

### Current Source Density Mapping

We used current source density (CSD) mapping to identify laminar boundaries in our recordings. While monkeys maintained fixation on the screen, 100% luminance contrast white annular stimuli were flashed for 32 ms, positioned at the center of the RF. The LFP signal following the stimulus onset was averaged across trials and spatially smoothed using a Gaussian kernel (sigma = 140 µm). The CSD was calculated as the second spatial derivative of the LFP:

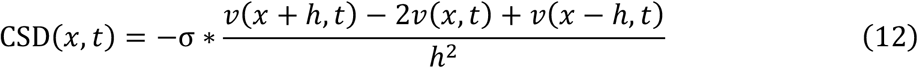

where *x* is the position in the extracellular medium at which the CSD is calculated, *t* the time following the stimulus onset (advancing in 1 ms increments), *h* the spacing between recording sites on the linear probe (here 70 μm), *v* the voltage, *σ* the conductivity of the cortical tissue (0.4 S/m). We interpolated the CSD every 7 μm. The input layer was identified by the boundaries of the early current sink characterizing feed-forward input into layer IV. Channels above and below this sink were classified as superficial and deep, respectively.

### Cued saccade task

During the task, subjects acquired and held fixation for a variable delay period (500–900 ms) prior to initiating a saccade in response to a target point appearing in the periphery. The simultaneous disappearance of the fixation point served as the go cue. After executing an accurate saccade, subjects then had to continue holding fixation at the target point for 500 ms to receive a reward. To prevent subjects from preemptively planning a saccade prior to the go cue, both the saccade target location and the delay period duration were pseudo-randomized. The target location was drawn from one of two possible locations, while the delay period duration was drawn from an exponential distribution. Targets were located 2.83 dva from the initial fixation point. Target locations were orthogonal to one another, and were each oriented 45° to and equidistant from the fixation to population receptive field axis. Only eye movements originating from <0.75 dva of the initial fixation point, and terminating <0.75 dva from the target were considered as successful trials. While the subjects executed these eye movements, oriented Gabor stimuli were continuously presented on a 13 × 13 grid (0.73 dva between adjacent grid points; Fig. 2B) spanning the visual region of interest at 60 Hz. The grid was centered on a point 4 dva from fixation along the fixation-receptive field axis.

Thus, the coordinates of saccade targets and population receptive fields within the frame of the stimulus grid were fixed (Fig. 2B). On each frame of stimulus presentation, a single stimulus drawn from one of 6 random orientations (0, 30, 60, 90, 120, 150°) was presented at a single grid location. Saccades were identified from eye-tracker data with a velocity-thresholding algorithm (68, 69). On average, subjects performed 895 trials of the cued saccade task (minimum of 729 trials, maximum of 1029 trials). Accumulating the recorded neuronal activity across trials provides comprehensive and dense coverage of all 13×13 grid locations for stimuli. Spiking activity from these trials was then combined by temporally aligning the data to saccade onsets, allowing to estimate the spatial receptive fields in a moment-by-moment manner relative to saccade onset.

### Data Analysis

Wherever possible, we analyzed our data using the estimation statistics framework (70, 71). The estimation statistics framework provides a principled way to measure effect sizes coupled with estimates of uncertainty, yielding interval estimates of uncertainty. To account for the temporal offset between the eye-tracker and the neural recording, an 8.33 ms offset has been applied to the data. This adjustment reflects the net difference between the estimated eye-tracker system latency of 16.66 ms (two frames at 120 Hz) and the mean photodiode offset of 8.33 ms.

### Continuous receptive field mapping

Receptive fields were mapped for each single unit as a function of time. Spikes were binned for each unit in response to each stimulus flash in a time window 50 to 100ms after flash onset. Stimulus flashes were then binned (41 ms centered window slid from -400 to 400ms relative to saccade onset by 1 ms) and their corresponding spike counts were averaged. This procedure generated a 13 × 13 grid of spike counts at each location for each timepoint relative to saccade onset, which was then linearly interpolated at tenfold spatial resolution and smoothed with a Gaussian kernel. Each timepoint was normalized such that the sum of all grid positions was equal to one to control for changes in firing rate. The location of receptive fields of individual units were determined by finding the center of mass of stimulus positions that elicited more than half the maximum firing. The receptive fields of individual neurons during fixation without saccade planning, termed ‘current receptive fields (CRFs)’, were calculated based on their responses to stimulus flashes presented during the pre-planning period. For each monkey, this period was defined from the latest time of the first stimulus flash relative to saccade onset (always preceded by fixation acquisition; -410.3 ms for monkey M, -423 ms for monkey D) to the earliest time of target appearance (-205.3 ms for monkey M, -211.4 ms for monkey D) across all trials. Since the visual cortex exhibited long-range correlated activity (72), neurons might respond to stimuli distant from their RFs. To avoid this potential effect, the CRF search was constrained to a 9 x 7 grid subarea (5.82 × 4.37 dva) of all possible flash positions (Fig. S2C). Starting from the appearance of the go cue, we constrained the RF search to a 4.37 × 4.37 dva subarea of all possible flash positions (Fig. S2B), which contained the CRFs (minimal eccentricity of 0.78 dva, maximal eccentricity of 2.40 dva) and the ideal remapped RFs (CRFs shifted by the cued saccade vectors) of all neurons which exhibited forward remapping. Only units with firing rates greater than 3 Hz were included in the analysis.

### Pre-saccadic suppression (PS)

For each neuron, the PS per saccade target location was computed based on the spiking responses to stimuli presented within the visual region of interest, which was a 9 x 8 grid subarea (5.82 × 5.09 dva; Fig. S2A):

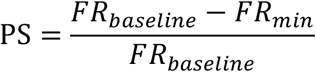

where *FR*_baseline_ was the baseline firing rate computed from the spike activity during the pre-planning period, defined as the interval before saccade target appearance, *FR*_m*in*_ the minimum firing rate during the reaction period, defined as the interval between target appearance and saccade onset (Fig. 2F). A positive PS indicates suppressed spiking activity, whereas a negative PS indicates enhanced neuronal activity.

### Comparison of neuronal responses to different planned saccade directions

For each neuron, the responses under different planned saccade directions were computed as the spiking responses to stimuli presented in the visual region of interest, which was a 9 x 8 grid subarea (5.82 × 5.09 dva; Fig. S2A). We computed a saccade direction dominance index (SDDI), which was calculated as:

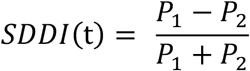

where P_1_ and P_2_ are the spiking responses under two saccade directions, respectively. By subsampling the response to stimuli presented at each location across trials, we obtained a distribution of SDDI and computed the 95% confidence interval of its mean. Neurons with a completely signed confidence interval were classified as saccade direction-biased neurons (Fig. 2E). In Fig. 2D, the responses and the associated SDDIs were calculated in a moment-by-moment manner as introduced in continuous receptive field remapping. In Fig. 2E, the SDDIs were computed using the spiking responses to stimuli presented during the period from -50 to 50 ms relative to saccade onset.

### Pairwise RF relationship analysis

For characterization of pairwise RF relationship, we only included neurons that exhibited predictive forward RF remapping by calculating for each neuron the distance between its temporal RFs and the ideal direction of forward remapping (the axis passing through the CRF along the saccade direction). Neurons with this distance constantly lower than a threshold value (1 dva for both monkeys in Fig. 3C, D) throughout the remapping period (from 50 ms before saccade onset to saccade offset) were identified as performing forward RF remapping. The preserved RF relationship held for a range of threshold values for this distance, which was larger for monkey M. We reasoned that it was because the CRF eccentricities of recorded units from monkey M were larger than that of monkey D (Fig. S2D), allowing for a larger threshold value to distinguish forward RF remapping from convergent RF remapping, where the RF was remapped to the saccade target.

The pairwise RF relationship was characterized by the pairwise difference of RF shifts, denoted as *Δremap* (Fig. 3B). To account for the variability in the locations of the stimulus grids across sessions, which were determined by the receptive fields of the recorded columns, the RFs of individual neurons were calculated within the frame of the stimulus grid, where the coordinates of the fixation point, the saccade targets, and thus the saccade directions were fixed. Only neurons with identical laminar identities were paired. To verify that the experimentally obtained *Δremap*s were significantly small, we compared them to the Δ*remaps* between neurons with shuffled CRFs. We generated the temporal evolution of RF shifts for 300 permutations of CRFs, from which the pairwise differences were calculated. The permutation of CRFs was constrained within neurons with identical laminar identities. The numbers of neurons from the superficial, input and deep layers included in the analysis (Fig. 3D) were: 28/48/24 under saccade target 1 and 26/42/20 under saccade target 2 for monkey M, 21/16/2 under saccade target 1 and 23/21/9 under saccade target 2 for monkey D.

The direction of *Δremap* was constrained to [0°, 180° ]. To account for the spontaneous noise in neural activity, the deformation of *Δremap* was calculated with reference to the mean *Δremap* during the pre-planning period and then compared with the saccade directions, which were 45°, 135° within the frame of stimulus grid (Fig. 3E, F).

### Characterization of stable error of remapping direction

To validate the model prediction that the error of remapping direction increases with CRF eccentricity, we characterized the stable remapped RF during saccade planning from the experimental data, which corresponded to the steady state of the network model under the input due to visual stimulus and CD input. For each neuron, we generated the temporal evolution of its average error of remapping direction across two saccade targets and calculated its temporal standard deviation, *std*_*RF*_(*t*), within a 30 ms window (advancing in 1 ms increments). The time of achieving stable remapping was defined as the earliest time when *std*_*RF*_(*t*) dropped below a certain threshold level, *θ*_*std*_, and persisted for a certain amount of time *T*_persist_. The average error of remapping direction over the 30 ms window centered at this time was taken as the estimated stable error. To focus on the period where remapping occurred and minimize the potential effects of realigned retina inputs after saccades, we only considered the time interval from when the temporal RF became closer to the ideal remapped RF (CRF shifted by the saccade vector) than to the CRF until saccade completion. In Fig. 5, *θ*_*std*_ = 0.3 rad, *T*_*persist*_ = 10 ms. The result held for a range of these hyperparameters.

