## Supplemental Material for "Pre-saccadic suppression and compensatory cortical distortions support predictive visual remapping"

*Monkey M*

| | Number of neurons involved in the analysis | Number of neurons that exhibit remapping | Ratio of remapping neurons |
|---|---|---|---|
| Superficial layer | 53 | 35 | 66.0% |
| Input layer | 79 | 58 | 73.4% |
| Deep layer | 46 | 29 | 63.0% |
| Total counts | 178 | 122 | 68.5% |

*Monkey D*

| | Number of neurons involved in the analysis | Number of neurons that exhibit remapping | Ratio of remapping neurons |
|---|---|---|---|
| Superficial layer | 91 | 37 | 40.7% |
| Input layer | 72 | 34 | 47.2% |
| Deep layer | 36 | 11 | 30.6% |
| Total counts | 199 | 82 | 41.2% |

*Both monkeys*

| | Number of neurons involved in the analysis | Number of neurons that exhibit remapping | Ratio of remapping neurons |
|---|---|---|---|
| Superficial layer | 144 | 72 | 50.0% |
| Input layer | 151 | 92 | 61.0% |
| Deep layer | 82 | 40 | 48.8% |
| Total counts | 377 | 204 | 54.1% |

**Table S1. Laminar distribution of analyzed V2 neurons**

Number of total neurons analyzed (restricted to single units with baseline firing rates greater than 3 Hz) and the corresponding number and proportion of neurons exhibiting significant forward RF remapping (see Methods) across cortical layers.

Figure S1

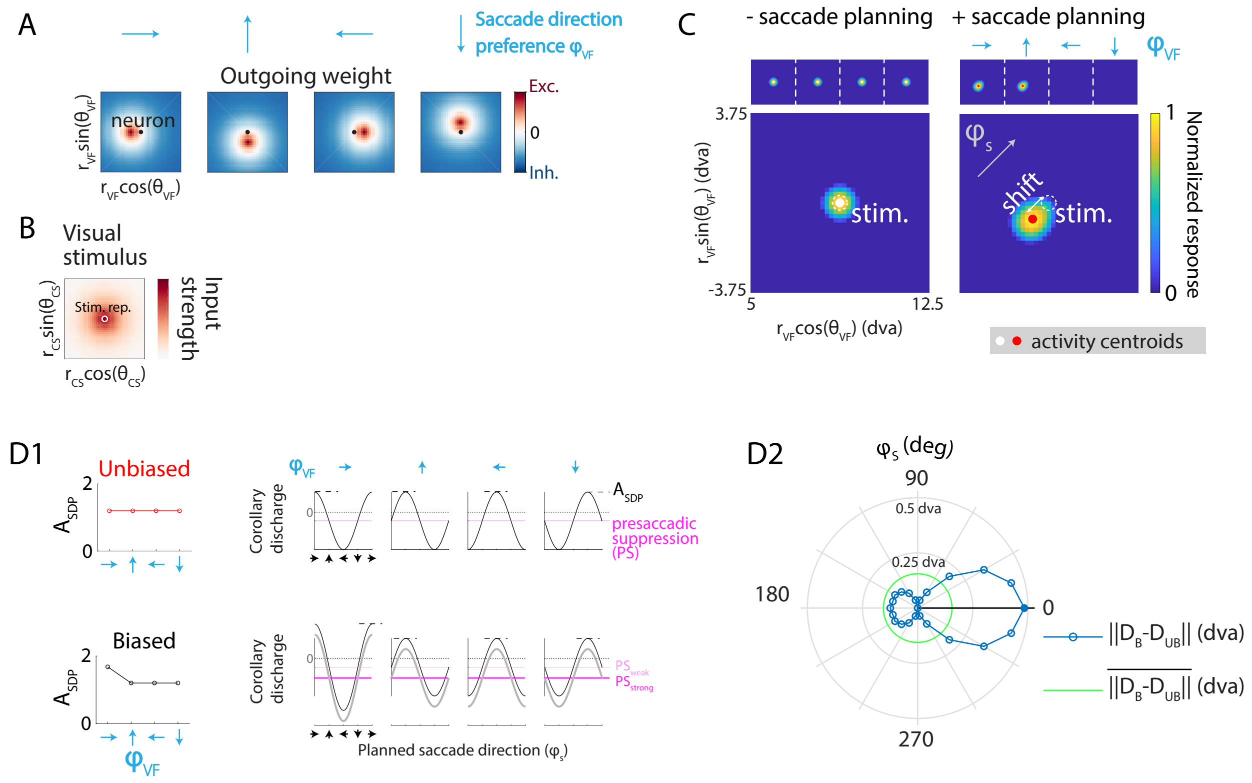


**Figure S1. Network model of receptive field (RF) remapping under uniform cortical transformation.**

(**A**) Outgoing weights from neurons with same spatial RFs (black dots) but four different SDPs (blue arrows), as a function of the RFs of postsynaptic neurons. (**B**) Amplitude of inputs caused by visual stimulus as a function of neurons' cortical locations. White circle: cortical representation of visual stimulus. (**C**) Steady-state population activity (by RF location) in response to a visual stimulus presented without (left) and with (right) saccade planning. Top row: activity grouped by saccade direction preferences (SDPs). Bottom row: activity averaged across SDPs. White dashed circles: location of the visual stimulus. Grey arrow: planned saccade direction ($\varphi_S$). (**D**) Remapping with biased SDP amplitude. **D1**: Left: Unbiased and biased SDP amplitude ($A_{SDP}$) as a function of SDP ($\varphi_{VF}$). Right: Corresponding corollary discharge amplitude for neurons with four different SDPs, as a function of planned saccade direction. **D2**: Remapping error induced by bias in the strength of saccade direction selectivity as a function of planned saccade direction. Black line: direction of SDP along which the bias is introduced. Green circle: mean error across tested saccadic directions ($\varphi_S$). Filled circle: maximum error across $\varphi_S$. $PS = 2.8$, $A_{SDP} = 1.68$ for $\varphi_{VF,i} = 0°$ under the biased condition and $1.2$ otherwise.

Figure S2

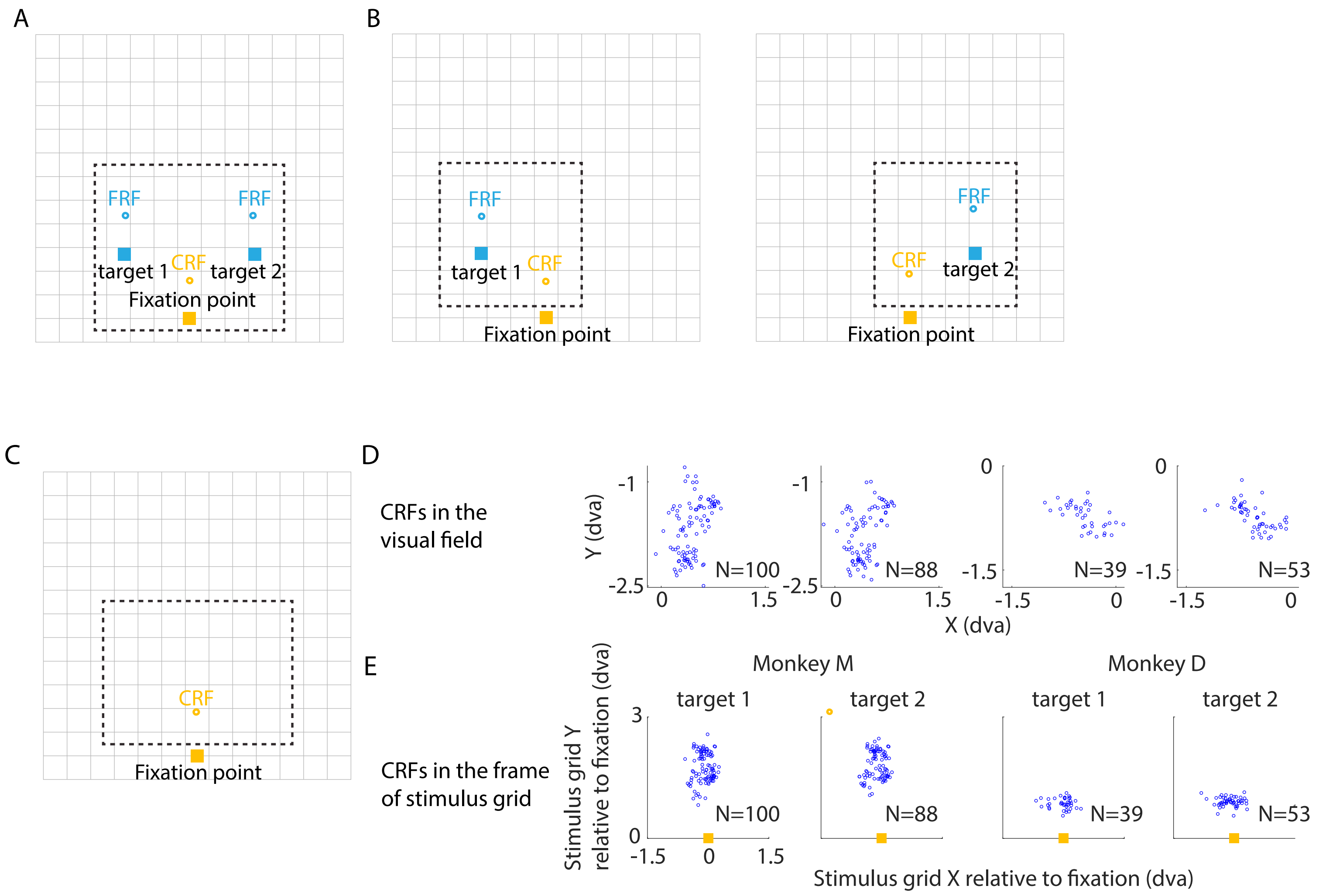


**Figure S2. Neuronal responses and RFs under a cued saccade task**

(**A**) The area of stimulus locations used to compute the responses under various saccade directions and characterize pre-saccadic suppression. (**B**) Visual regions of interest for RF searching during saccade planning for various saccade directions. (**C**) Visual region of interest for RF searching during the pre-planning period. (**D**) Distribution of current RFs (CRFs) for recorded neurons exhibiting forward remapping under various saccade directions for each monkey. (**E**) Same as (D) but shown in the reference frame of the stimulus grid, where the coordinates of fixation point and saccade targets were fixed across sessions.

Figure S3

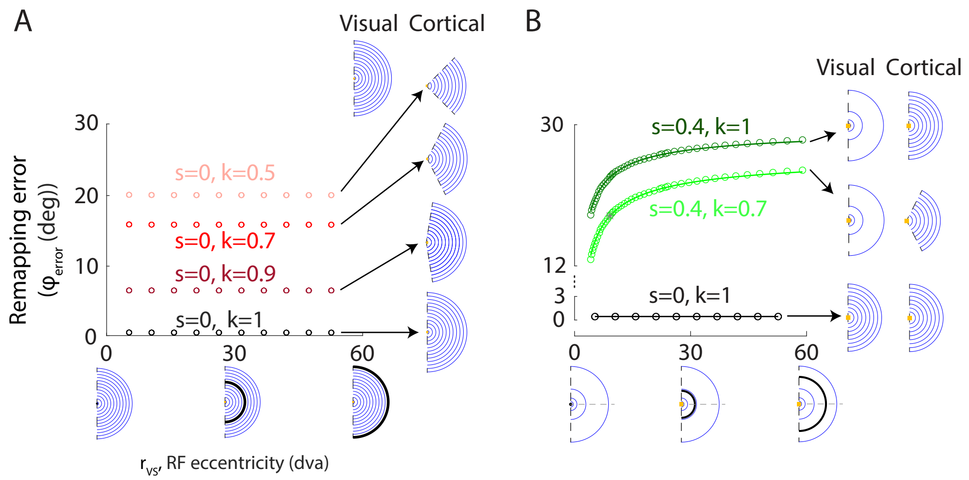


**Figure S3. Remapping error under polar angle contraction with eccentricity-independent cortical magnification**

(A) Same as Fig. 4D but under conditions of eccentricity-independent cortical magnification (CM) ($s = 0$) across various extents of polar angle contraction ($k$). The remapping error is independent of CRF eccentricity. (B) Light green: same as Fig. 4D but under conditions of both eccentricity-dependent CM ($s = 0.4$) and polar angle contraction ($k = 0.7$). Black: uniform transformation as shown in Fig. 4D. Dark green: eccentricity-dependent cortical magnification (CM) only as shown in Fig. 4D. Grey asterisk: transformations considered in Fig. 4B, C. Curves: fits using general power functions (see Methods).

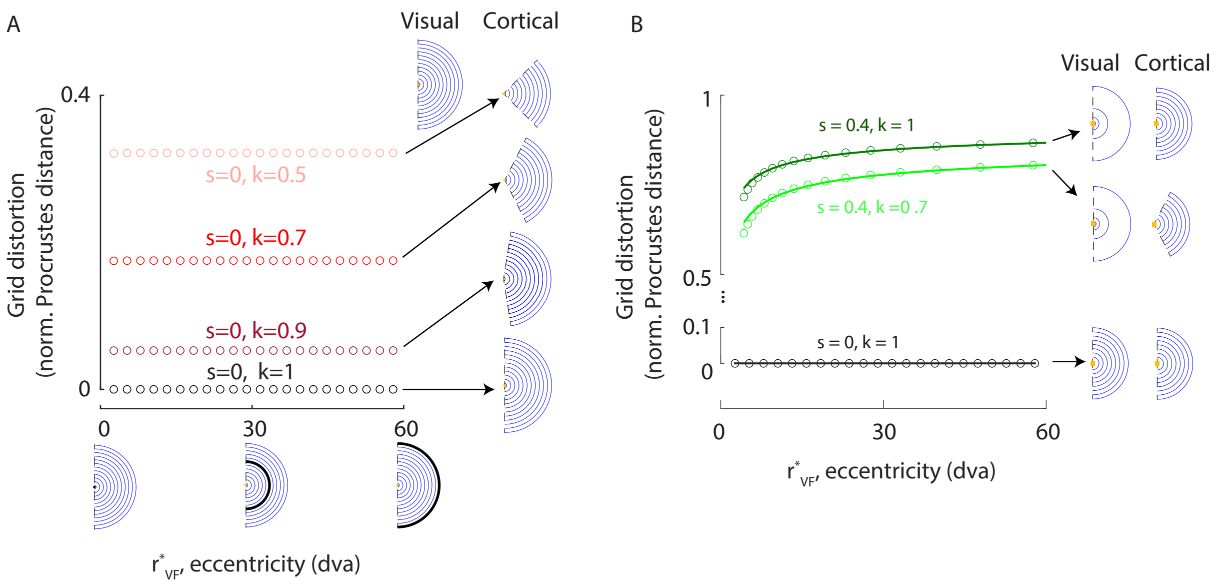


**Figure S4. Grid distortion under polar angle contraction with eccentricity-independent cortical magnification**

(A) Same as Fig. 6B but under conditions of eccentricity-independent CM ($s = 0$) across various extents of polar angle contraction ($k$). The grid distortion is independent of the eccentricity of network location. (B) Light green: same as Fig. 6B but under conditions of both eccentricity-dependent CM ($s = 0.4$) and polar angle contraction ($k = 0.7$). Black: uniform transformation as shown in Fig. 6B. Dark green: eccentricity-dependent cortical magnification (CM) only as shown in Fig.6B.

Figure S5

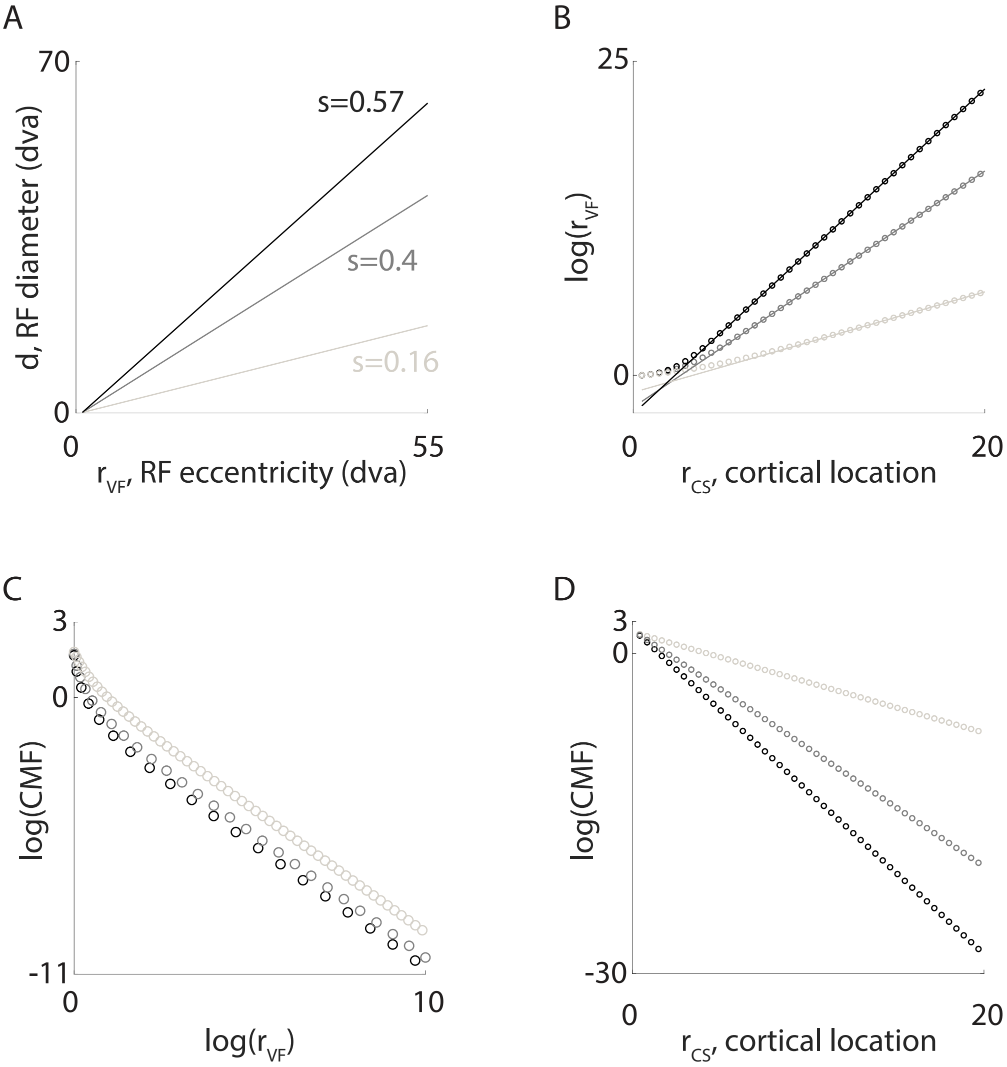


**Figure S5. Constructing eccentricity-dependent cortical magnification**

(**A**) Linear relationships between RF diameter ($d$) and RF eccentricity ($r_{VF}$) explored in the study. (**B**) Corresponding $log(r_{VF})$ as a function of cortical location ($r_{CS}$), constructed using the linear relationship shown in (A). Circles: constructed data. Lines: theoretical linear relationships (Eq. 13-14). (**C**) Corresponding CM factor (CMF), approximated by forward difference methods based on (**B**). (**D**) Same as (C) but as a function of $r_{CS}$. $D_c = 0.5$, $d_1 = 0.1$, $\mathrm{r}_{\mathrm{VF,1}} = 1$.
